# Probe-assisted Structural Topography Reveals In-tissue Lesion Heterogeneity in Alzheimer’s Disease

**DOI:** 10.64898/2026.11.17.767196

**Authors:** Fuzerong Wang, Xiaoting Sun, Qinyue Zhao, Yue Hu, Lu Xu, Shen Zhang, Bingyao Cui, Dan Li, Jing Zhang, Wentao Chen, Cong Liu, Kiryl D. Piatkevich, Dan Yang

## Abstract

Disease lesions are typically defined by morphology, yet this classification often masks underlying structural heterogeneity. In Alzheimer’s disease (AD), amyloid-β (Aβ) plaques appear as featureless deposits under conventional microscopy, obscuring distinct sub-regions with divergent packing, maturation, and pathological potential. Here, we present Probe-Assisted Structural Topography of Amyloid (PASTA), a single-staining framework that combines a novel pan-Aβ probe (NapIM) with fluorescence lifetime-resolved detection and super-resolution microscopy to unmask this hidden complexity. Leveraging PASTA, we quantify plaque maturity via the Protofibril Index (PFI) at sub-lesion level, and resolve the three-dimensional (3D) in-tissue architecture of Aβ plaques at ∼30 nm resolution, identifying radially extended Aβ branches as fundamental structural units. PASTA enables structural mapping across plaque developmental stages to reconstruct their dynamic growth trajectories, resolving two distinct assembly lineages (dense and loose) with divergent growth paths that converge to a mixed terminal subtype and share a common transition point. We further applied PASTA for mapping microglial engagement, linking interdigitated microglial contacts to dense plaque compaction and uncontacted fibril branches to loose morphology. Together, PASTA provides an unprecedented framework for understanding Aβ heterogeneity, nanoscale architecture, and microglial-mediated remodeling in AD.

## Introduction

Morphological appearance has long served as the primary criterion for defining disease lesions^1–3^. Lesions such as amyloid-β (Aβ) plaques in neurodegenerative disorders, which appear homogeneous or indistinguishable under conventional microscopy, are often categorized into broad, uniform groups^4–8^. However, this apparent uniformity frequently masks inherent structural and molecular heterogeneity. Emerging evidence indicates that what appears as a solid, featureless deposit may actually consist of distinct sub-regions with varying Aβ distribution^9–11^ and maturity^12–18^.

Critically, recent cryo-electron tomography (cryo-ET) studies have revealed that Aβ plaques are composed of highly ordered, parallel bundles of Aβ fibrils^19, 20^—a finding that suggests fundamental structural uniformity. This raises two key questions: how these uniform building blocks assemble into the heterogeneous architecture observed in tissues, and whether this heterogeneity encodes a dynamic history of plaque formation.

Existing imaging approaches address only partial aspects of this problem. Fibril-specific dyes (e.g., thioflavin S) label dense plaque cores but fail to detect protofibrillar regions; antibodies cannot penetrate compact aggregates; and super-resolution techniques relying on antibody labeling are constrained by the same penetration limitations. To date, no single method has achieved molecularly specific, heterogeneity-resolved, and three-dimensionally (3D) super-resolved mapping of the entire Aβ plaque landscape within intact tissue. The inability to resolve these fine-scale architectural details—due to the diffraction limit of light and the lack of molecularly specific contrast agents—has impeded our understanding of how these structures assemble, evolve, and interact with their microenvironment. Thus, there is an urgent need for imaging strategies that can unmix this hidden information and reveal the nanoscale architecture of pathological assemblies in their native tissue context.

Here, we present probe-assisted structural topography of amyloid (PASTA), an integrated approach that combines a highly specific pan-Aβ probe (NapIM) with fluorescence lifetime imaging microscopy (FLIM) and expansion microscopy (ExM). This method enables molecularly specific, heterogeneity-unmixed, and 3D super-resolved mapping of Aβ plaque architecture and their cellular interactions in Alzheimer’s disease (AD).

Using PASTA, we introduce the Protofibrillar Index (PFI), a quantitative metric that converts fluorescence lifetime values into a continuous readout of Aβ maturity. This metric reveals plaque sub-regions and rare subtypes that are invisible to intensity-based imaging. Integrating PASTA, we resolved the 3D nanoscale architecture of plaques and identified radially arranged Aβ branches as a fundamental structural unit. Systematic PASTA mapping of a plaque cohort in the hippocampus across progressive AD stages allowed us to identify two distinct assembly lineages (dense and loose), reconstruct their dynamic growth trajectories, and uncover a mixed terminal subtype arising from the convergence of both lineages.

We further demonstrate that microglial engagement drives the compaction of dense-type plaques via interdigitated contacts, whereas branches escaping microglial coverage retain a loose-type morphology. Collectively, PASTA provides a versatile platform for elucidating the structural basis of amyloid pathology, offering unprecedented insights into plaque evolution and microglial function that advance our understanding of plaque heterogeneity and cellular dynamics in AD.

### Design and characterization of PASTA

To enable heterogeneity-resolved mapping of plaque architecture, we developed a pan-Aβ fluorescent probe, NapIM, which exhibits fluorescent responses to Aβ at different aggregation states and displays state-discriminative fluorescence lifetimes (Fig. 1a–c; Extended Data Fig. 1). This differentiating capability stems from its sensitivity to viscosity and polarity (Extended Data Fig. 1) ^21^. NapIM efficiently binds to both off- and on-pathway Aβ species across various aggregation states (Extended Data Fig. 1). To validate NapIM’s binding mechanism to Aβ fibrils, we incubated brain-extracted Aβ fibrils (from the postmortem brain of a single AD patient) with the probe to form NapIM-bound Aβ fibrils (NapIM-Aβ complex, Type I Aβ fibril; Extended Data Fig. 2a–d). Cryoelectron microscopy (cryo-EM) was used to obtain electron density maps of NapIM-Aβ and apo-Aβ fibrils at an overall resolution of 3.3 Å (Fig. 1d,e; Supplementary Table 1). Compared to apo-Aβ fibrils, the addition of NapIM caused negligible changes to the helical parameters or 2D class average patterns of Aβ fibrils (Extended Data Fig. 2e; Supplementary Table 1). The complex showed two additional strong densities corresponding to NapIM within a deep pocket at the head-to-tail interface of the two protofilaments (Fig. 1d,e; Extended Data Fig. 2f). Reconstruction was performed based on a 4-fold rise (as a single NapIM molecule spans four amyloid layers), and NapIM was successfully fitted into the density maps with high fidelity (Fig. 1f).

**Fig. 1:**
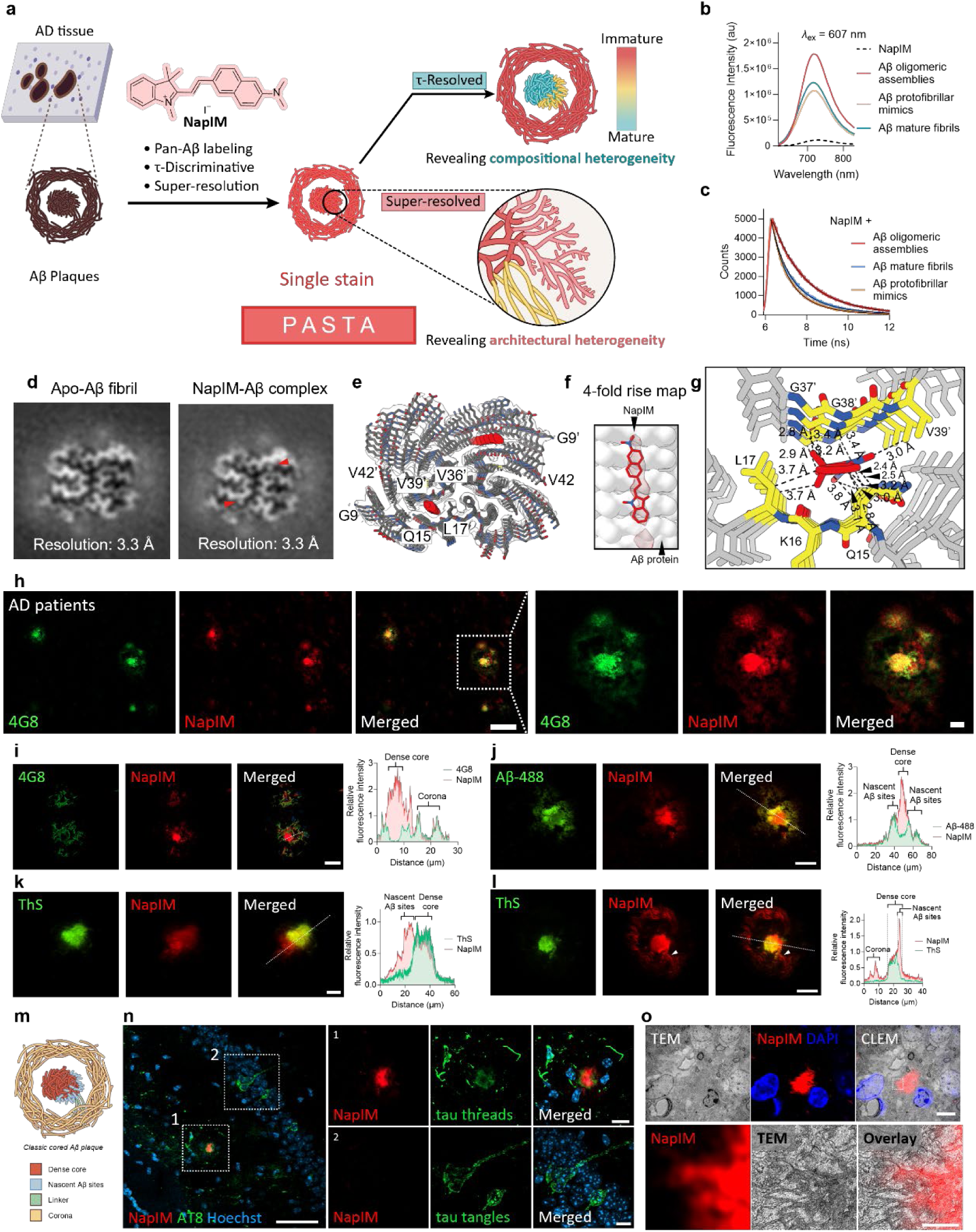
Design and characterization of PASTA. **a**, A schematic of PASTA to discriminate plaque heterogeneity and achieve nanoscale plaque architecture. The chemical structure of NapIM is shown. **b**, Fluorescent spectra of NapIM (10 µM in PBS, pH 7.4) in the presence of Aβ42 oligomeric assemblies, protofibrillar mimics, and mature fibrils (20 µM each). **c**, Fluorescence decay profiles of NapIM (10 µM in PBS, pH 7.4) in the presence of oligomeric assemblies, protofibrillar mimics, and mature fibrils (20 µM each) determined with excitation at 635 nm. Fluorescence lifetimes, Aβ oligomeric assemblies: τ = 1.83 ± 0.02 ns; Aβ protofibrillar mimics: τ = 1.17 ± 0.01 ns; Aβ mature fibrils: τ = 1.40 ± 0.01 ns. **d**, Central slices of cryo-EM 3D maps of the type I fibril with and without the addition of NapIM. Red arrows indicate the extra densities. The corresponding resolutions of each map were labeled below. **e**, Structural model of NapIM-Aβ complex fitted in the density map. The density map is restricted to the area within a 2-Å radius of the structural model. Extra densities are colored red. The residues forming the NapIM-binding hole are labeled. **f**, Side view of the density maps is shown with a 4-fold helical rise repeat distance. The fitted chemical molecule is displayed, and the Aβ protein model is shown in a grey surface. **g**, Ligands are shown in red, and interacting residues are highlighted in yellow. Interactions between the ligands and Aβ holes are indicated, with interacting residues and distances (Å) labeled. **h**, Representative confocal images of Aβ plaques labeled by NapIM and Aβ/APP antibody (4G8) from post-mortem brains of individuals with AD. Right, magnified view of plaque region (boxes in left panel). Scale bars, 50 µm (left), 10 µm (right). **i–l**, Representative magnified confocal images of Aβ plaques labeled by NapIM, Aβ/APP antibody (4G8) (**i**), fluorescently labeled Aβ42 (**j**), and ThS (**k,** compact plaque, and **l,** classic cored plaque) from 13-month-old (**i,j,l**) or 9-month-old (**l**) 5×FAD mice. Scale bars, 20 µm (**i**, **j**), 10 µm (**k**, **l**). Right, line profiles of fluorescence intensities across the plaques that are shown in the left panels. The linker between the dense core and corona is indicated by white arrows in **l**. **m**, Schematic of classic cored Aβ plaque depicting dense core, protofibrillar hotspots, corona, and linker connecting dense core and corona. **n**, Representative confocal images of NapIM and phospho-Tau (labeled by AT8) from a 21-month-old 3×Tg mouse. Right, magnified views of Aβ plaque, tau threads, and tau tangles (boxes in left panel) showing NapIM selectively labels Aβ plaque over tau materials. Scale bars, 50 µm (left); 10 µm (right). **o**, Top, representative correlative light and electron microscopy (CLEM) images of Aβ plaques from a 9-month-old 5×FAD mouse showing that NapIM staining patterns colocalized with amyloid protein compositions identified by TEM. Bottom, magnified view of NapIM-labeled Aβ plaque (white box in top panel) with corresponding region in TEM. Scale bars, 5 µm (top); 2 µm (bottom).

Models constructed for both NapIM-Aβ and apo-Aβ fibrils revealed that the probe binding pocket is formed by residues Q15–L17 (from one protofilament) and V36–V39 (from the opposing protofilament; Fig. 1e–g; Extended Data Fig. 2g), with a binding mode similar to that of AV-45—a clinically approved FDA PET tracer for Aβ fibrils (Extended Data Fig. 2h) ^22^. The NapIM-Aβ complex model revealed several key interactions: hydrophobic interactions between NapIM and the main chains of K16 (from one protofilament) and G38 (from the opposing protofilament, denoted G38’), as well as hydrophobic interactions between NapIM and the side chains of Q15 and V39’ (Fig. 1g). Overall, the atomic structure of the NapIM-Aβ complex is consistent with results from fluorescent spectroscopy, confirming NapIM’s binding to Aβ fibrils. The size-restricted hydrophobic pocket selectively accommodates the insertion of the hydrophobic NapIM molecule, enabling specific interaction with Aβ fibrils (Extended Data Fig. 2g).

To fully verify NapIM’s specificity for amyloid deposits in the complex biological context, a double fluorescent staining assay was performed using brain sections from the postmortem brain of an AD patient with an Aβ-specific antibody. This assay showed that NapIM specifically stained dense and focal Aβ plaques (Fig. 1h), confirming its specific binding to Aβ species in AD tissue. To further characterize the probe’s selectivity and specificity, we tested its performance in imaging Aβ plaques in the 5×FAD animal model—an AD model with aggressive amyloid pathology that facilitates the easy acquisition of freshly harvested brain tissue for various subsequent assessments^23^.

NapIM enabled high-contrast visualization of Aβ plaques with minimal non-specific labeling (Extended Data Fig. 3a), and its in-tissue emission peak matched that observed in in-tube studies (Extended Data Fig. 3b,c). Pan-Aβ staining in brain tissue via NapIM was confirmed by co-staining with antibodies on fixed tissue (Fig. 1i; Extended Data Fig. 3d–i), fluorescently labeled Aβ peptide (Aβ-488) labeling on fresh tissues (Fig. 1j) ^24^, and co-staining with well-established probes (Fig. 1k,l; Extended Data Fig. 4a–d) ^24–27^. NapIM efficiently labeled all subregions of Aβ plaques, including dense cores, coronas, and nascent sites—outperforming conventional Aβ antibodies in labeling protofibril-rich regions (coronas and nascent sites) (Fig. 1i,m; Extended Data Fig. 3g–i) and traditional probes in labeling mature fibril-rich regions (dense cores) (Fig. 1k–m; Extended Data Fig. 4d) ^15, 24^.

In brain slices from 3×Tg mice, NapIM exhibited no fluorescence at AT8-positive Tau fibrillary threads or tangles (Fig. 1n; Extended Data Fig. 4e), confirming its high specificity for Aβ over Tau. This outperforms traditional probes such as ThS and Methoxy-X04, which also bind to Tau aggregates in tissue staining^20, 28^. Combining transmission electron microscopy (TEM) with NapIM staining, we further observed that the structural features of plaques visualized via TEM matched NapIM’s staining pattern—including branches at plaque edges and amyloid fibril bundles visible in high-resolution TEM—confirming that NapIM staining delineates amyloid-related features in plaques (Fig. 1o; Extended Data Fig. 4f,g).

In summary, we identified NapIM as a specific Aβ probe that efficiently labels all subregions of Aβ plaques. Based on these findings, we employed this probe to map the structural topography of densely packed Aβ plaques via a single-staining approach—a strategy we term probe-assisted structural topography of amyloid (PASTA).

### PASTA enables discrimination of Aβ plaque heterogeneity and maturity

Pathological brain samples are typically derived from fixed tissues, which inherently limit the application of live-cell molecular assays (as shown in Fig. 1j). To overcome this limitation, we leveraged PASTA to rapidly and efficiently unveil hidden information within these fixed specimens via single staining. By integrating with fluorescence lifetime imaging microscopy (FLIM) ^29^, PASTA enables rapid discrimination of distinct subregions and subtypes of plaques in fixed brain tissue (Fig. 2a), extracting new dimensions from previous information-limited fluorescence signals. The PASTA channel selectively exhibits longer fluorescence lifetimes (τ) in protofibril-rich regions and shorter τ values at mature fibril-rich sites (Fig. 2b).

**Fig. 2:**
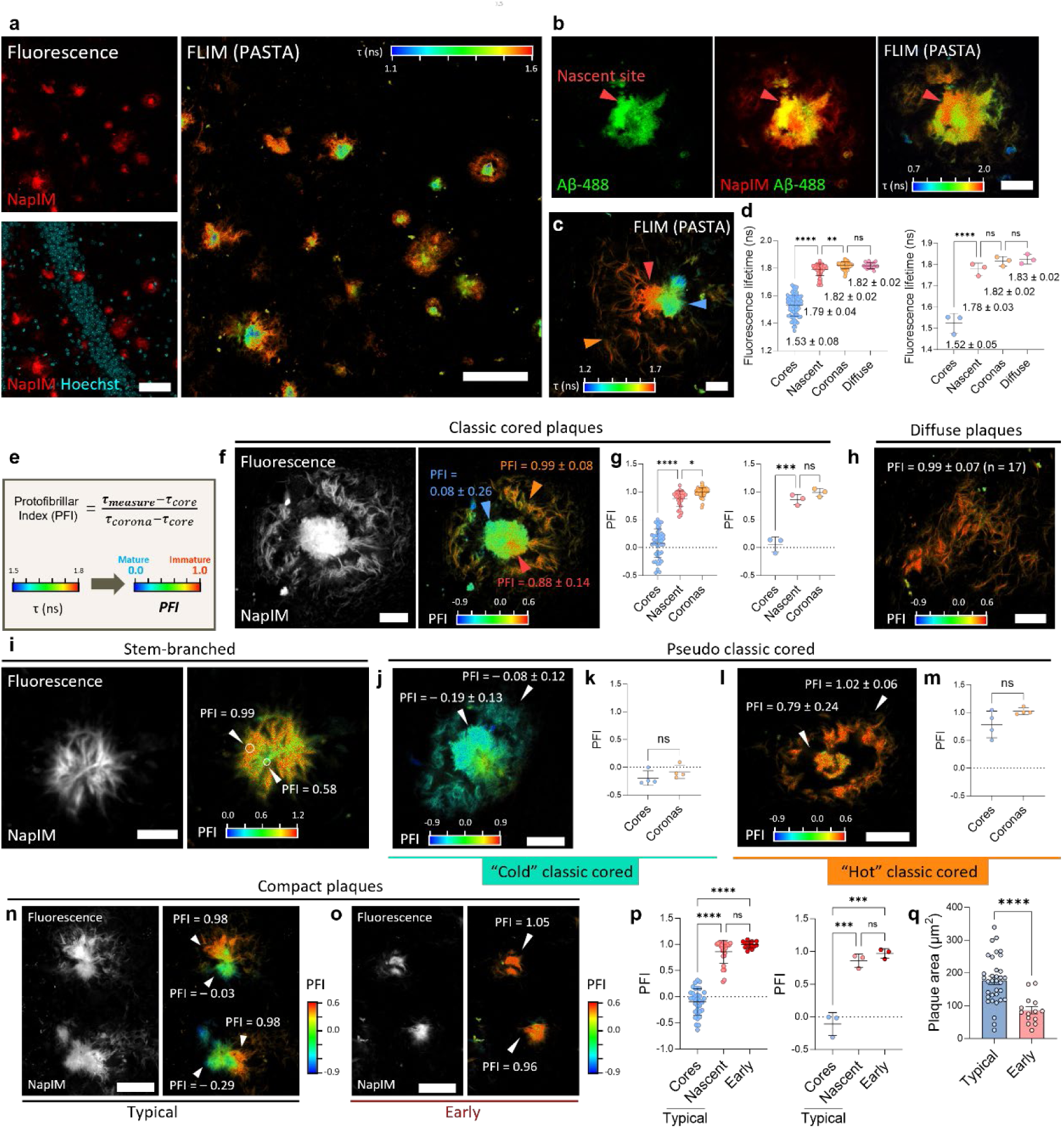
Discrimination of Aβ plaque heterogeneity and maturity by PASTA. **a**, Representative fluorescence (left) and lifetime (right) images of Aβ plaques from a 12-month-old 5×FAD mouse labeled by NapIM. Scale bars, 50 µm. **b**, Representative fluorescence (left, middle) and lifetime (right, NapIM signal) images of an Aβ plaque from a 12-month-old 5×FAD mouse labeled by NapIM and fluorescently labeled Aβ42-488 to show nascent sites. Scale bar, 10 µm. **c**, A representative lifetime (NapIM signal) image of an Aβ plaque containing core, corona, and nascent sites from a 12-month-old 5×FAD mouse labeled by NapIM. Nascent site, corona, and core are indicated by red, orange, and blue arrows, respectively. Scale bar, 10 µm. **d**, Fluorescence lifetime values of Aβ plaque subregions, shown by either individual plaques (left) or mice (right). *n* = 95 plaques from *n* = 3 mice (12-month-old, 5×FAD). Fluorescence lifetime values (mean ± s.d.) of corresponding subregions or plaque type are indicated. **e**, Conversion of fluorescence lifetime values to protofibrillar index (PFI). **f**, Representative fluorescence (left, middle) and PFI (right) images of a classic cored plaque from a 12-month-old 5×FAD mouse labeled by NapIM. Scale bar, 10 µm. **g**, PFI values of classic cored plaque subregions, shown by either individual plaques (left) or mice (right). *n* = 44 classic cored plaques from *n* = 3 mice (12-month-old, 5×FAD). **h**, A representative PFI image of a diffuse plaque from a 12-month-old 5×FAD mouse labeled by NapIM. Scale bar, 10 µm. **i**, Representative fluorescence (left) and PFI (right) images of a stem-branched Aβ plaque from a 12-month-old 5×FAD mouse labeled by NapIM. Scale bar, 5 µm. **j**, A representative PFI image of a “cold” classic cored plaque from a 12-month-old 5×FAD mouse labeled by NapIM. Scale bar, 10 µm. **k**, PFI values of “cold” classic cored plaque subregions. *n* = 4 “cold” classic cored plaques from *n* = 3 mice (12-month-old, 5×FAD). **l**, A representative PFI image of a “hot” classic cored plaque from a 12-month-old 5×FAD mouse labeled by NapIM. Scale bar, 10 µm. **m**, PFI values of “hot” classic cored plaque subregions. *n* = 4 “hot” classic cored plaques from *n* = 3 mice (12-month-old, 5×FAD). **n–o**, Representative fluorescence (left, middle) and PFI (right) images of typical compact plaques (**n**) and early compact plaques (**o**) from a 12-month-old 5×FAD mouse labeled by NapIM. Scale bars, 20 µm. **p**, PFI values of compact plaque subregions and subtypes, shown by either individual plaques (left) or mice (right). *n* = 34 typical compact plaques and *n* = 14 early compact plaques from *n* = 3 mice (12-month-old, 5×FAD). **q**, Quantification of the plaque areas in typical or early compact plaques. *n* = 34 typical compact plaques and *n* = 14 early compact plaques from *n* = 3 mice (12-month-old, 5×FAD). \*\*\*\**P* < 0.0001 by unpaired Student’s *t*-test. **f**, **h**, **j**, **l**, PFI values (mean ± s.d.) of corresponding subregions or plaque type are indicated. Statistical analysis was performed using one-way ANOVA for multiple comparisons (**d**, **g**, and **p**) and two-tailed unpaired t-tests (**k**, **m**, and **q**). Data are mean ± s.d.

Leveraging this contrast, we aimed to utilize PASTA-derived τ values to quantitatively distinguish plaque subtypes (cored and diffusive) and, within cored plaques, to resolve specific subregions (cores, nascent sites, and coronas). Unlike the corona, which surrounds the core and can be directly distinguished by morphological features, nascent sites and dense cores are tightly adjacent, rendering them difficult to resolve via single-staining approaches. The PASTA channel enables robust discrimination between nascent sites and dense cores, exhibiting significantly higher τ values in nascent sites (1.789 ± 0.043 ns) than in dense cores (1.530 ± 0.077 ns) (Fig. 2b–d).

Through morphological analysis, we identified coronas and diffuse plaques, which exhibited comparable PASTA-derived τ values, suggesting that these regions share similar micro-compositions (Fig. 2c,d). Nascent sites, coronas, and diffuse plaques all exhibited similarly higher τ values than dense cores, indicating underlying similarities among protofibril-rich domains (nascent sites, coronas, and diffuse plaques) (Fig. 2c,d). We observed that certain plaques exhibit transitional τ values at subregion interfaces (Fig. 2c; Extended Data Fig. 5a), suggesting that PASTA-derived τ values can effectively reflect gradients in micro-composition. Building on this observation, we propose a *Protofibrillar Index* (PFI) based on these τ values to quantify the relative abundance of protofibrils (Fig. 2e). In this metric, values approaching 1 indicate a composition similar to the immature, protofibril-rich corona, whereas values nearing 0 indicate a mature, fibril-rich nature comparable to the dense core.

This conversion from PASTA-derived τ to the PFI scale requires the determination of reference τ values for the corona and core within the specific sample. We observed that in classic cored plaques, corona regions consistently exhibited PFI values approaching 1, while dense cores yielded values near 0, characterized by exceptionally low standard deviations (Fig. 2f). This minimal variability establishes classic cored plaques as reliable intrinsic standards for calibration. Practically, minimal sampling (n = 3 plaques per mouse) is sufficient to define the PFI scale. Notably, PFI values derived from this limited sampling showed no significant difference compared to the mean calculated from the entire dataset (Extended Data Fig. 5b–e).

Diffuse plagues were defined as structures with a loose, fibrous morphology and PFI values approaching 1 (Fig. 2h). Their lack of distinct subregions suggests a homogeneously protofibril-rich composition (Fig. 2h). In contrast, ThS (a fibril-specific probe) exhibited uniformly distributed lifetimes across all plaque regions, providing no information to distinguish between subregions or plaque types (Extended Data Fig. 5f). A primary application of PASTA-derived PFI values is to investigate rare and ambiguous plaque types. We identified a “stem-branched” Aβ plaque morphology, characterized by a clear main stem and side branches—resembling a feather structure. The main stem showed a PFI of approximately 0.6, while the side branches exhibited values around 1 (Fig. 2i), suggesting the side branches are more protofibril-rich and potentially undergoing active growth alongside the main stem.

Interestingly, we also identified two categories of pseudo-classic cored plaques that are indistinguishable from normal classic cored plaques when relying solely on morphological features visualized via fluorescence intensity (Fig. 2j,l). Unlike normal classic cored plaques, these plaques exhibited homogeneous PFI values across their entire structure: “cold” classic cored plaques display a corona with PFI approaching 0, indicating protofibril-poor coronas, whereas “hot” classic cored plaques exhibit a core with PFI near 1, indicating protofibril-rich cores (Fig. 2j–m). These findings underscore that relying exclusively on fluorescence intensity and morphology can lead to the misclassification of rare or anomalous plaques as common subtypes, resulting in misinterpretations of true plaque heterogeneity.

Compact plaques are traditionally defined as dense deposits featuring radiating materials but lacking distinct coronas. Under the PASTA modality, typical compact plaques exhibit pronounced zonal heterogeneity (Fig. 2n). Surprisingly, PASTA uncovered a distinct subset within this category, which we term early compact plaques (Fig. 2o). Unlike their typical counterparts, these plaques displayed a homogeneous distribution of PFI values approaching 1 throughout the entire structure, lacking any observable zonal differentiation (Fig. 2n–p). This uniformity suggests that the plaque is in a state of active, synchronous growth. Consistent with this hypothesis, the mean area of early compact plaques was significantly smaller than that of typical compact plaques (Fig. 2q).

In summary, by leveraging PASTA-derived τ values and the PFI, we achieved the discrimination of both plaque subtypes and subregions via a single-staining approach while preserving high imaging resolution. This strategy provides a method for the unbiased quantitative classification of plaques and the rigorous analysis of plaque heterogeneity.

### Mapping of Aβ dense cores at a single-branch resolution by PASTA

PASTA-revealed plaque heterogeneity suggests that structures appearing similar under conventional confocal microscopy may be intrinsically distinct. In contrast, conventional confocal microscopy often depicts Aβ dense cores as featureless, solid deposits due to limited resolution, thereby hindering the detailed mapping of their hidden structural diversity. To visualize the fine architecture of plaques more clearly, we integrated PASTA with protein retention expansion microscopy (ExM) ^31^. ExM embeds tissues in isotropically swellable hydrogels that physically separate densely packed biomolecules and magnify sub-diffraction structures upon hydration, thereby enhancing resolution proportional to the linear expansion factor.

Compared with five conventional Aβ probes, NapIM proved most suitable for ExM, delivering superior signal contrast (CBR > 9.5) compared to the others (CBRs < 1.5) (Fig. 3a,b; Extended Data Fig. 6a,b). By integrating PASTA with ExM, we achieved significant resolution enhancement of Aβ plaques with minimal distortion (Fig. 3c,d; Extended Data Fig. 6c). Leveraging this super-resolution PASTA modality, we resolved a typical radial arrangement of strand-like structures within the plaques (Fig. 3e–g), which we term “Aβ branches.” These Aβ branches, while densely packed side-by-side, exhibit distinct inter-branch spacing (Fig. 3e–g; Extended Data Fig. 6d). The PASTA channel further showed that the termini of some Aβ branches diverge, creating a dendritic morphology at the plaque periphery (Fig. 3g, blue arrows). Additionally, intense sprouts along the branches formed a feather-like morphology (Fig. 3h–i), with short, rod-like barbs (approximately 70 nm in diameter) projecting from branches that were approximately 200 nm in diameter (Fig. 3i,j).

**Fig. 3:**
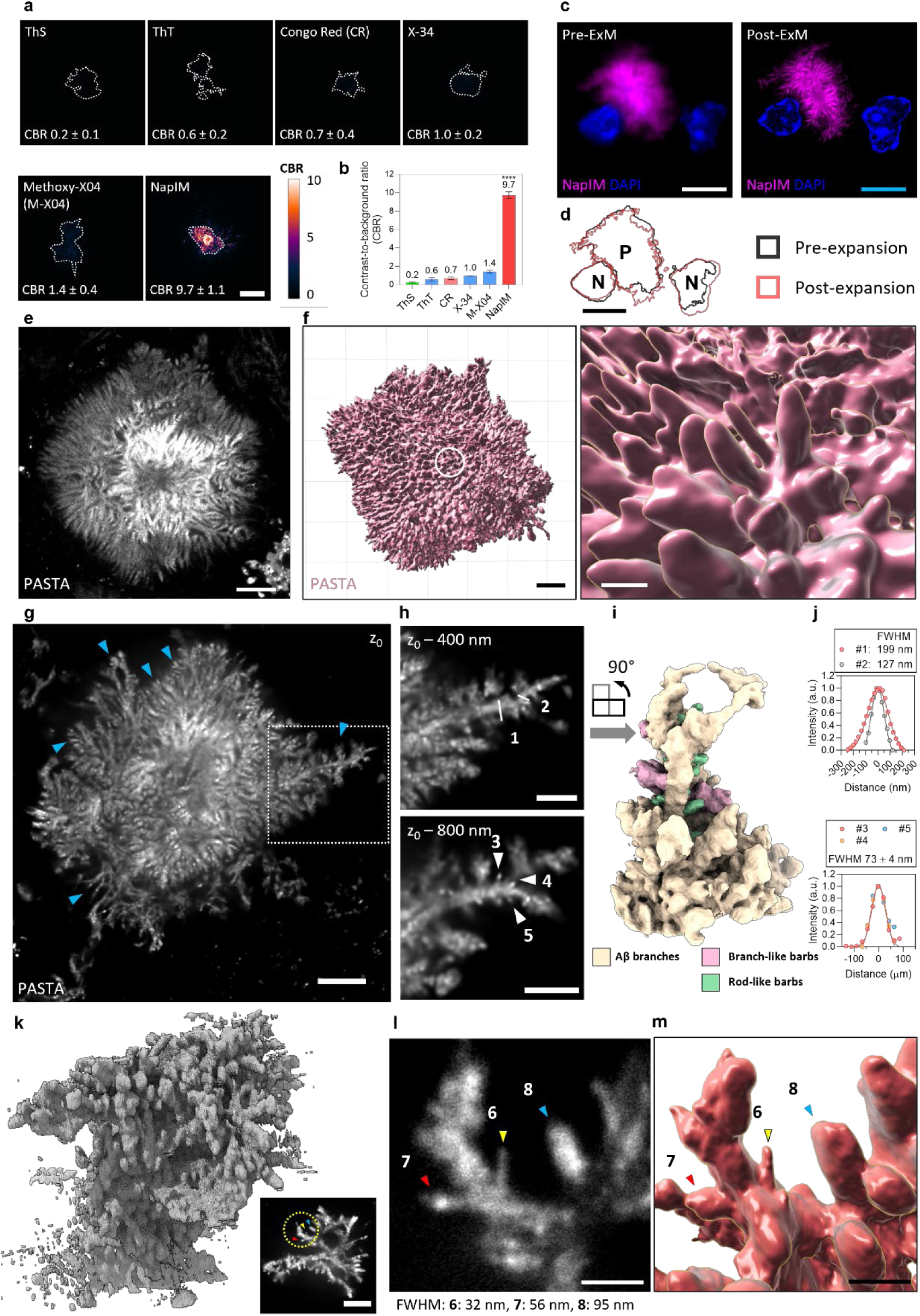
Nanoscale branching architecture visualized by PASTA with ExM. **a**, Comparison of NapIM and other Aβ probes (ThS, ThT, Congo Red, X-34, and Methoxy-X04) as Aβ staining for ExM. The color bar indicates the contrast to background ratio (CBR) ^30^ of labeling. The average CBR of each probe (mean ± s.d., *n* = 9 regions) is indicated. Brain slices were collected from the same 12-month-old 5×FAD mouse and expanded 5.1-fold. Scale bar, 10 µm (biological scale, equals to physical scale/expansion factor). **b**, Quantifications of the CBRs for the images shown in **a**. Error bars, s.e.m., *n* = 9 regions. The average values were indicated above the column. \*\*\*\**P* < 0.0001, NapIM *vs* the other group, by one-way ANOVA for multiple comparisons. CR, Congo Red. M-X04, Methoxy-X04. **c**, Representative pre- and post-expansion (5.6-fold) images of an Aβ plaque from a 3-month-old 5×FAD mouse labeled by NapIM to show the enhanced resolution. Scale bars, 5 µm for physical scale (white) and biological scale (light blue). **d**, The binary mask of plaques and nuclei in (**c**). Scale bar, 5 µm. P, Aβ plaques. N, Nuclei. **e**, A representative ExM image (6.9-fold) of a spikey Aβ plaque from a 22-month-old 5×FAD mouse labeled by NapIM. Scale bar, 2 µm (biological scale). **f**, Left, 3D reconstruction from z-stack images from **e** to show the 3D structure of Aβ plaque. Right, magnified region as indicated by the circle in **f** to show distinct Aβ branches. Scale bars, 2 µm (left) and 1 µm (right) in biological scale. **g**, A representative ExM image (4.5-fold) of an Aβ plaque from a 12-month-old 5×FAD mouse labeled by NapIM. Blue arrows indicate representative branching sites. Scale bar, 2 µm (biological scale). **h**, Magnified regions indicated by the dashed box in **g** at different z-sections. Scale bar, 1 µm (biological scale). **i**, 3D reconstruction from z-stack images from the dashed box in **g** to show the 3D structure of Aβ barbs. **j**, Gaussian fit of the fluorescence profiles of signal intensity in **h** marked by white lines (**h,** upper) or white arrows (**h,** below) with numbering to yield FWHM. Symbols, experimental data; lines, Gaussian fit of the individual profiles. **k**, 3D reconstruction from z-stack images of a small plaque (inserted panel), revealing its complete spatial architecture. Inserted panel, A representative ExM image (6.5-fold) of an early Aβ plaque from a 12-month-old 5×FAD mouse labeled by NapIM. Scale bar, 1 µm (biological scale). Three typical branches (labeled 6, 7 and 8) are indicated by arrows. **l**, Magnified region as indicated by the dashed circle in **k**. FWHM values of 3 branches (labeled 6, 7 and 8) are indicated below the image. **m**, 3D reconstruction from z-stack images from **l** to show the 3D structure of Aβ branches. Scale bars, 1 µm (biological scale).

To better visualize the 3D orientation and morphology of plaques, we analyzed an early-stage plaque characterized by its smaller size and lower compactness (Fig. 3k). The 3D reconstruction revealed intricate structural complexity with numerous branch-like protrusions extending from the core (Fig. 3k). This 3D reconstruction further confirmed the presence of rod-like barbs protruding from Aβ branches, with clear endpoints and diameters as small as 32 nm (Fig. 3l,m; Extended Data Fig. 6e–h). Structured illumination microscopy (SIM) imaging on non-expanded tissues showed analogous branch-like structures, thereby independently confirming the existence of this architecture (Extended Data Fig. 6i–l).

Together, these data establish Aβ branches as the fundamental structural unit of plaques, featuring a radially organized, barb-bearing architecture previously invisible to light microscopy. This finding provides a direct structural link between nanoscale Aβ fibrils and the tissue-level plaque.

### Characterization of Plaque Phenotypes and Heterogeneity Driven by PASTA

Driven by PASTA’s robust capacity for super-resolution mapping, we characterized the diverse phenotypes and intrinsic heterogeneity of plaques. We applied PASTA to map plaques in the hippocampal regions of brain sections from 3-month-old 5×FAD mice, which uniquely captures the full continuum of plaque maturation from nascent to advanced states. Leveraging PASTA’s superior imaging capabilities—specifically its high contrast and resolution—we performed unbiased sampling across a large cohort of plaques, enabling comprehensive classification. Surprisingly, we identified striking heterogeneity in plaque assembly patterns, in addition to variations in size (Fig. 4a). Notably, this heterogeneity extends to the fine topology of individual Aβ branches (Fig. 4b). We observed one subset of branches with prominent spikey sprouts, creating a rough, dense appearance, whereas another subset exhibited smooth, loose contours entirely devoid of such protrusions (Fig. 4b). This distinct variation in Aβ branch morphology prompted us to investigate the underlying architectural assembly modes governing plaque structures.

**Fig. 4:**
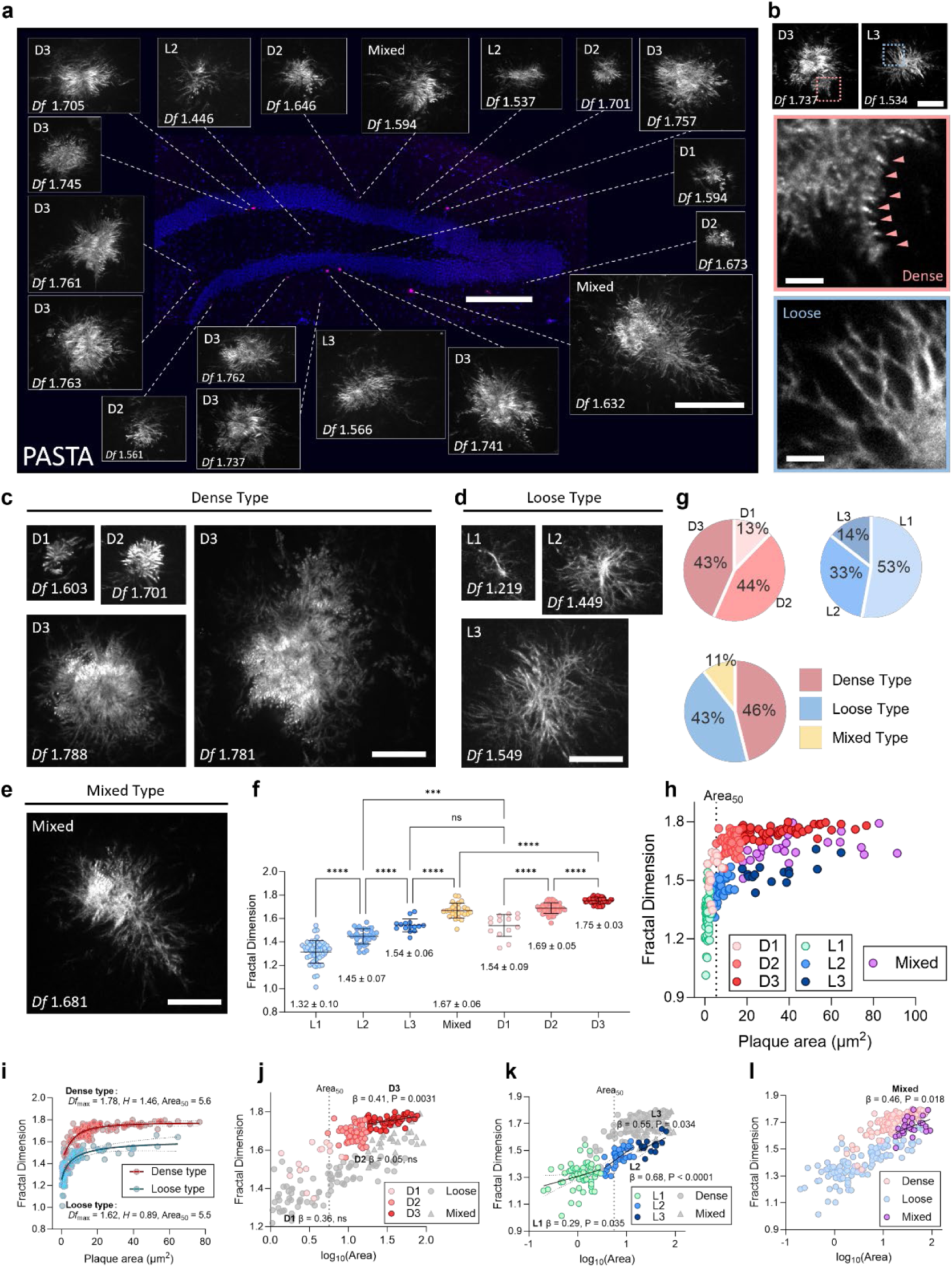
Aβ plaque phenotyping and evolutionary trajectories by PASTA. **a**, A representative fluorescence map of a DG region from a 3-month-old 5×FAD mouse staining by NapIM (magenta) and Hoechst (blue), with each plaque further visualized to show nanoscale architecture. Scale bars, 200 µm (map) and 10 µm (PASTA images) in biological scale. **b**, Top, representative fluorescence images showing branches of dense (left) and loose (right) appearances. Bottom, magnified regions indicated by the colored dashed boxes in the top panels. Scale bars, 5 µm (top), 1 µm (bottom) in biological scale. **c–e**, Representative fluorescence images of dense type plaques (**c**) with stages D1, D2, D3, loose type plaques (**d**) with stages L1, L2, L3, and mixed type plaques (**e**), each showing 3 representative images. All images are shown at the same scale for ease of comparison, with all scale bars of 5 μm in biological scale. **f**, Fractal dimension (*D_f_*) of plaque subtypes and stages in a cohort of *n* = 246 plaques from *n* = 3 mice (3-month-old, 5×FAD). **g**, The proportion of each stage in *n* = 118 dense type plaques (top left), *n* = 105 loose type plaques (top right) from *n* = 3 mice (3-month-old, 5×FAD), and the proportion of dense, loose, and mixed in total *n* = 246 plaques (bottom) from *n* = 3 mice (3-month-old, 5×FAD). Plotting *D_f_* versus plaque areas shows the distribution of each subtype and stage. **h–l**, Scatterplots showing the correlation between *D_f_* and plaque areas of dense type (**i, j**), loose type (**i, k**), and mixed type (**l**). Nonlinear (**i**) and linear (**j–l**) trend lines are shown, and the 95% confidence intervals are indicated by dashed lines. *D_f,max_*, maximum fractal dimension. *H*, Hillslope: *H* > 1 signifies positive cooperativity yielding a sharp, “switch-like” transition, while *H* < 1 reflects negative cooperativity or population heterogeneity resulting in a gradual response. Area*_50_*, half-maximal area. Coefficients of determination (R^2^) are 0.65 (dense type, **i**), 0.60 (loose type, **i**), 0.17 (D3), 0.08 (L1), 0.46 (L2), 0.30 (L3), and 0.21 (mixed type). The estimated regression coefficients (β) and P values are shown. ns, not significant (P > 0.05). In all plaque images, plaque types and stages (D1, D2, D3, L1, L2, L3, and mixed type) and *D_f_* values are indicated.

Through extensive mapping, analysis, sorting, and classification of a plaque cohort (n = 246 hippocampal plaques from n = 3 mice), we identified three distinct assembly patterns. The first, termed the “dense type”, is characterized by tightly packed, radially arranged Aβ branches with narrow inter-branch spaces, resulting in a compact appearance (Fig. 4c). The second, designated the “loose type”, features loosely arranged radial branches with wide inter-branch gaps, producing a vine-like, sprawling morphology (Fig. 4d). Building on these two categories, we identified a third “mixed type”, in which regions exhibiting both dense-type and loose-type characteristics coexist within a single plaque (Fig. 4e).

Interestingly, our analysis revealed that both lineages follow a size-dependent three-stage progression yet exhibit divergent structural trajectories: the dense type evolves through tightly packed stages (D1: 13%, D2: 44%, D3: 43%), whereas the loose type progresses as loosely arranged forms (L1: 53%, L2: 33%, L3: 14%) (Fig. 4c–d,g; Extended Data Fig. 7a; classification criteria summarized in the Methods section). In contrast, the mixed type (11% of total, vs. 46% dense and 43% loose) presents a unique profile, characterized almost exclusively by large dimensions (minFeret > 8 µm), indicative of an advanced evolutionary stage (Fig. 4e,g; Extended Data Fig. 7a). Crucially, these morphological distinctions are underpinned by fundamental differences in fractal geometry. The fractal dimension (*D_f_*) quantifies how completely a pattern fills space, distinguishing between simple, smooth geometries and highly branched, space-filling architectures. Quantitative analysis of the *D_f_* of plaque images revealed significant disparities between the loose and dense populations, providing numerical validation for our morphological classification (Fig. 4f). Importantly, such quantitative fractal analysis is only feasible because PASTA resolves the intricate internal architecture of plaques. Rendering plaques as featureless solid masses via conventional imaging would preclude meaningful *D_f_* characterization.

The loose type exhibited significantly lower *D_f_* values compared to the dense type (Fig. 4f), with the mixed type bridging the gap between these two extremes. A *D_f_* value approaching 1 for the loose type indicates a dominance of linear, lower-dimensional characteristics, suggesting a growth strategy prioritizing sprawling extension over area occupation (Fig. 4f). In contrast, the dense type showed *D_f_* values approaching 2, reflecting a tendency to fill the surrounding space, maximizing area coverage rather than linear reach (Fig. 4f). Furthermore, the mixed type displayed intermediate *D_f_* values situated between those of L3 and D3, consistent with its hybrid structural nature (Fig. 4f).

Consistently, scatter plot analysis (*D_f_* versus plaque area) revealed distinct trimodal clustering of plaque structural features (Fig. 4h; Extended Data Fig. 7b–d). Within their respective lineages, the early stages (L1 and D1) clustered in the exponential growth phase, characterized by rapid fluctuations in *D_f_* values, whereas the mature stages (L3 and D3) occupied the plateau phase, marked by slowed changes in *D_f_* values—indicating structural maturation (Fig. 4h; Extended Data Fig. 7b–d). The intermediate stages (L2 and D2) were distributed within the transition zones bridging L1/D1 and L3/D3 (Fig. 4h; Extended Data Fig. 7b–d). Notably, the mixed-type plaques formed a distinct cluster situated precisely between the plateau phases of the L3 and D3 populations (Fig. 4h,l; Extended Data Fig. 7d).

The maximum fractal dimension (*D_f,max_*) values for the pure lineages were distinct—1.78 for the dense type and 1.62 for the loose type—indicating that independent maturation of either lineage does not spontaneously generate mixed characteristics (Fig. 4i). This quantitative gap strongly supports the intrinsic independence of the mixed type as a unique entity. Both dense and loose lineages exhibited a positive correlation between plaque area and *D_f_* values, suggesting that as Aβ content increases, plaques progressively enhance their space-filling capacity (Fig. 4i–l). Intriguingly, the scatter distribution of the mixed type closely adjoined the cluster of L2 and D2 (Fig. 4h; Extended Data Fig. 7d), suggesting a potential divergent developmental trajectory in which L2 and D2 plaques may transition into mixed-type architecture.

Notably, the dense type displayed threshold-dependent explosive growth (Fig. 4i, Dense type Hillslope = 1.46), reflecting a switch to rapid compaction during plaque evolution, while the loose type displayed a gradual accumulation profile (Fig. 4i, Loose type Hillslope = 0.89). These distinct kinetic signatures align with the spikey, highly branched architecture of the dense type versus the smooth, elongating filaments of the loose type. Despite these distinct growth dynamics, similar critical areas in both dense and loose lineages (Fig. 4i) highlight a potential universal window for intervention for both plaque phenotypes.

In summary, by integrating nanoscale morphological details with quantitative fractal analysis, PASTA has uncovered hidden structural heterogeneity and reconstructed the dynamic evolutionary landscape of Aβ plaques. Crucially, the superior resolution of PASTA renders these complex architectures amenable to rigorous mathematical description, revealing distinct growth modes previously obscured by conventional imaging and enabling a shift from morphological classification to objective phenotyping.

### Microglia contacts define plaque assembly modes

Having established the distinct structural phenotypes of amyloid plaques (dense, loose, and mixed types), we next sought to investigate the mechanisms driving these divergent assembly patterns. Aβ plaques reside within a complex cellular milieu characterized by microglial clustering, astrogliosis, and dystrophic neurite formation ^24, 32–35^—all of which could potentially modulate plaque architecture. Consistent with prior reports ^26, 33, 34, 36–39^, our imaging revealed that microglial processes are intimately associated with Aβ plaques (Extended Data Fig. 8a). Microglia either enveloped large regions of the plaque periphery or formed discrete pockets around individual Aβ branches (Fig. 5a, arrows c1–c3), and the gap between the dense core and corona of classic cored plaques was largely occupied by microglial cell bodies (Extended Data Fig. 8b)—suggesting that microglial interactions may shape plaque architecture. We also identified Aβ branches extending beyond the microglial barrier with little to no cellular coverage (Fig. 5a, arrow e1), consistent with observations by Condello et al. ^24^.

**Fig. 5:**
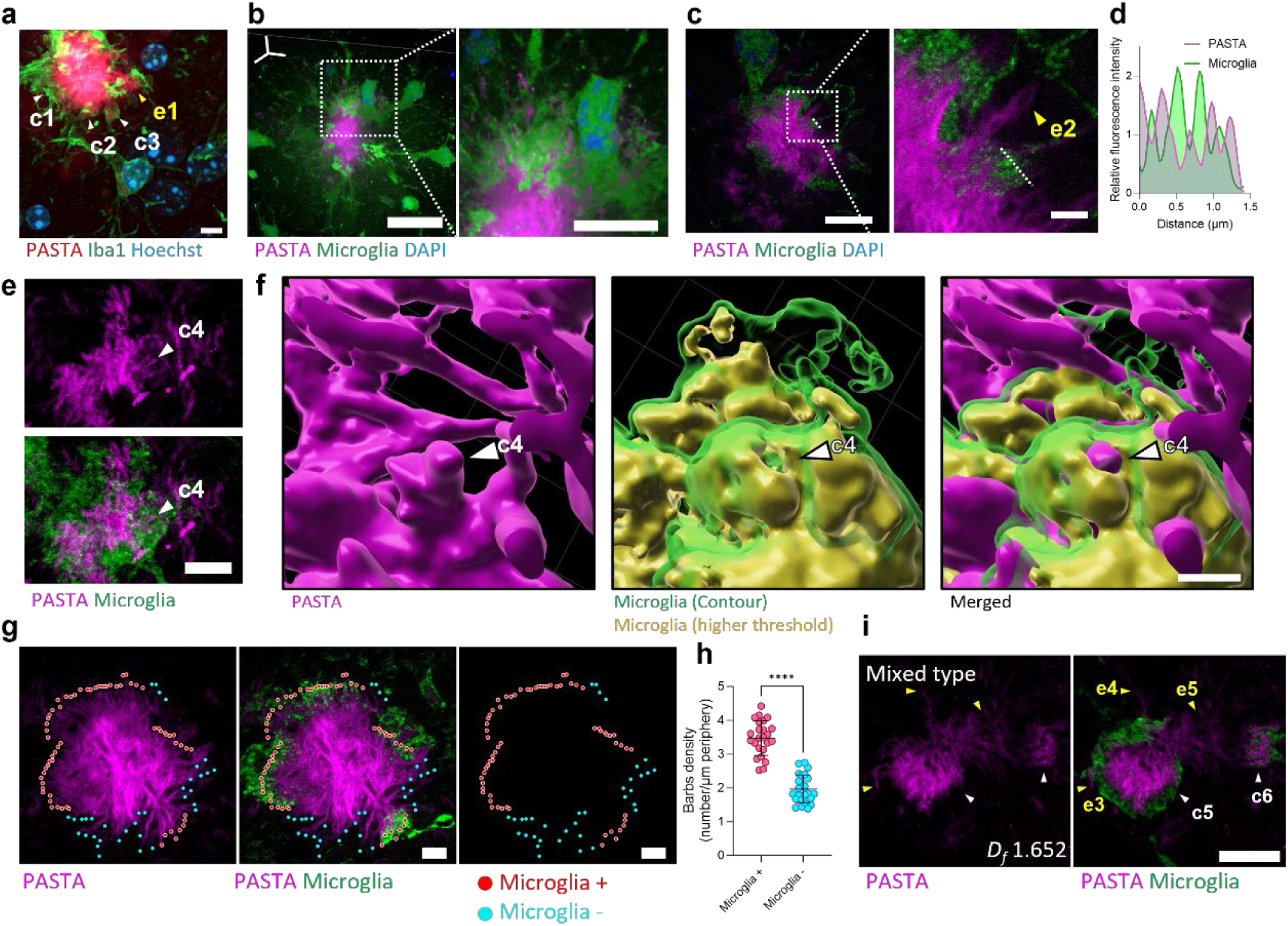
Visualization of microglial interaction with Aβ plaques. **a**, Representative magnified confocal images of Aβ plaques from a 10-month-old 5×FAD mouse labeled by NapIM and microglia-specific antibody (Iba1). Microglial covering and pockets were indicated by white arrows (c, covering), and escaping Aβ branches were indicated by yellow arrows (e, escaping). Scale bar, 5 µm. **b**, A representative 3D ExM image (6.5-fold) of Aβ plaque (labeled by NapIM) and microglia by z-stacking. Scale bar, 10 µm. Right, Magnified region of the dashed box. Scale bar, 5 µm (biological scale). **c**, Representative ExM images (6.0-fold) of Aβ plaques (labeled by NapIM) and microglia. Scale bar, 5 µm (biological scale). Right, magnified region of the dashed box. Scale bar, 1 µm (biological scale). **d**, Line profile of fluorescence intensities across the interaction sites as shown by the dashed line in **c**. **e**, Representative ExM images (5.5-fold) of Aβ plaques (labeled by NapIM) and microglia showing Aβ branches encapsulated by microglia. **f**, 3D reconstruction from z-stack images from **e** to show the 3D structure of Aβ fiber and microglial encapsulation. Scale bars, 2 µm (biological scale). The white arrows in **e** and **f** indicate the same barb. **g**, Representative ExM images (5.5-fold) of Aβ plaques (labeled by NapIM) and microglia showing higher Aβ barbs density under microglia encapsulation. Scale bar, 2 µm (biological scale). **h**, Quantification of the number of Aβ barbs per µm periphery (in biological scale) of the plaque (number/µm periphery) at microglia + or − regions. *n* = 24 plaques. \*\*\*\**P* < 0.0001 by unpaired Student’s *t*-test. **i**, Representative ExM images (4.5-fold) of Aβ plaques (labeled by NapIM) and microglia showing escaping Aβ branches. Microglial covering was indicated by white arrows, and escaping Aβ branches were indicated by yellow arrows. Scale bar, 5 µm (biological scale). **c–i**, Microglia were labeled by crossbreeding with CX3CR1-EGFP mice. Scale bars are in biological scale for ExM images (**c–i**). Microglia-covered branches are labeled as c1-c6. Escaping branches are labeled as e1-e5. All Cx3cr1-2A-EGFP-labeled tissue slices were collected from 9-month-old Cx3cr1-2A-EGFP/5×FAD +/+ mice.

Next, we utilized Cx3cr1-2A-EGFP/5×FAD +/+ mice to yield uniform microglial signals in thick tissues (Extended Data Fig. 8c) without antibody labeling. Microglia were found to extend filamentous processes at contact sites with Aβ plaques (Extended Data Fig. 8d–g), indicating intimate Aβ-branch-level interactions. Building on these observations, we employed super-resolved PASTA mapping and confirmed tight contacts between microglial processes and plaque peripheries (Fig. 5b). Notably, the diverging termini of Aβ branches were tightly enveloped by microglial processes, creating interdigitated fluorescent signals in cross-sectional profiles (Fig. 5c,d; Extended Data Fig. 9a–c). 3D reconstruction of the PASTA channel further revealed that microglia fully encase individual Aβ sprouts from a branch (Fig. 5e,f, arrow c4; Extended Data Fig. 9d). Larger, mature plaques were often partially covered, leaving exposed regions (Fig. 5c,e,g,i; Extended Data Fig. 9b,c), whereas smaller, early-stage plaques exhibited more extensive coverage, with some being fully encapsulated (Extended Data Fig. 9e).

Crucially, the structural morphology of Aβ branches appeared to be directly modulated by the presence of microglia. Regions of Aβ plaques in contact with microglia exhibited a significantly higher density of barbs, adopting a “denser” appearance compared to the “looser” morphology of microglia-free regions (Fig. 5g,h). This effect was evident even at the single-branch level: individual Aβ branches partially covered by microglia displayed a marked increase in barb density on the contacted side versus the exposed side (Extended Data Fig. 9f,g), suggesting that microglial contact promotes barb formation and structural compaction.

Aβ branches that “escape” microglial coverage (Fig. 5c, arrow e2; Fig. 5i, arrows e3–e5) exhibited a loose-type filamentous morphology with few to no sprouts—sharply contrasting with the spikey morphology of microglia-covered Aβ branches (Fig. 5g; Fig. 5i, arrows c5–c6; Extended Data Fig. 9f,g). This difference implies that microglial interaction is a key driver in the formation of dense-type characteristics and that the absence of such interaction allows branches to retain or adopt a loose configuration. Consequently, the coexistence of covered and uncovered regions within a single plaque may contribute to the formation of mixed-type architecture.

Finally, PASTA imaging of classic cored plaques confirmed that escaping branches directly connect to the surrounding coronas (Extended Data Fig. 9h, 10a), with coronas consisting of branchless, loose-type Aβ structures (Extended Data Fig. 9f–h, 10a). This suggests that the corona represents an extension of the loose-type growth mode, potentially formed by Aβ branches that have evaded microglial containment—thereby sharing a common compositional and conformational origin.

## Discussion

Disease tissues are often viewed as static endpoints, a constraint particularly critical in neurodegenerative disorders like Alzheimer’s disease (AD), where human brain samples are exclusively available post-mortem. Deciphering the complex developmental history embedded within these lesion snapshots is therefore essential to understanding disease progression. The accumulation of β-amyloid (Aβ) into plaques is a key pathological feature of AD ^40^. Conventional confocal microscopy depicts Aβ plaques as featureless, solid deposits, but this apparent uniformity masks profound underlying heterogeneity across three dimensions: gross morphology, composition, and nanoscale architecture. In human brain tissue, gross morphological assessment alone has identified at least seven distinct plaque types^4, 7, 8^—still the sole dimension resolvable by standard optical imaging. Relying exclusively on gross morphology yields limited insight, raising critical questions: Do morphologically distinct plaques share underlying micro-similarities? Can we move beyond subjective visual classification to establish objective quantitative criteria for plaques?

Techniques such as infrared (IR) imaging^12, 13, 15^ and Raman imaging^14–18^ have revealed significant compositional heterogeneity beneath homogeneous deposits, highlighting regional variations in plaque maturity. However, their utility is constrained by limited spatial resolution, low signal specificity, and challenges in distinguishing specific signals from background noise. To address this, we adopted a strategy of spectral and spatial deconvolution: PASTA enables high-resolution quantitative deconvolution of plaque compositional heterogeneity through PASTA-derived τ values and the introduction of the Protofibrillar Index (PFI). In our model, a PFI approaching 1 signifies an immature, protofibril-rich state, whereas a PFI approaching 0 indicates a mature, fibril-dense state. Our analysis reveals that within a single mature plaque, distinct domains coexist: typically, a central mature core (PFI ≈ 0) surrounded by multiple immature peripheral regions (PFI ≈ 1).

Remarkably, despite vast morphological differences, diverse plaque types exhibit similarities in their PFI profiles. The dense core of classic cored plaques, the compact ends of typical compact plaques, and other mature regions all converge at PFI ≈ 0. Conversely, the coronas and nascent sites of classic cored plaques, the nascent ends of compact plaques, diffuse plaques, and early compact plaques all show PFI ≈ 1—indicating shared mature and immature compositional signatures across these varied structures. Furthermore, our quantitative approach uncovered previously indistinguishable subgroups (e.g., “hot” versus “cold” classic cored plaques), which differ significantly in their maturity profiles. These findings underscore the necessity of transitioning from subjective morphological classification to objective, quantitative deconvolution to fully understand AD’s pathological landscape.

Revealing nanoscale architectural heterogeneity requires imaging capabilities that transcend the diffraction limit. While super-resolution techniques (STED, SMLM) can resolve sub-diffraction structures, antibody-based approaches are limited by poor penetration into dense cores ^41–43^, and small-molecule SMLM only images plaque peripheries ^44^. By contrast, PASTA identified Aβ branches: ∼200 nm filamentous units radially arranged within plaques, with dendritic termini and 30–70 nm barbs projecting from their surfaces (Fig. 3). This radial organization mirrors “amyloid stars” previously observed only by electron microscopy (EM) ^32–34^, but PASTA offers faster, large-volumetric mapping with higher molecular specificity—critical for linking structure to dynamic processes. Given that individual Aβ fibrils are 4–12 nm in diameter ^45–47^, the 200 nm Aβ branches likely form via fibril bundling, consistent with cryo-ET data ^19, 20^—establishing Aβ branches as the fundamental structural unit of plaques and resolving the longstanding question of how Aβ fibers assemble into tissue-level pathology.

Leveraging this spatial resolution, we performed comprehensive geometric and fractal analyses across mapped plaques, revealing a clear developmental progression: plaques initiate as either loose-type or dense-type, proceeding through early (L1, D1), intermediate (L2, D2), and late-stage (L3, D3) phases (Extended Data Fig. 10b). Importantly, both lineages converge at a similar critical area threshold marking the transition from early to intermediate phases—confirming this transition as an intrinsic structural threshold rather than a subjective classification. Our modeling further identifies distinct growth dynamics: the dense-type, characterized by high branching propensity, exhibits rapid expansion, while the loose-type, with limited branching, follows a gradual accumulation profile (Extended Data Fig. 10b). These lineages may exert different mechanical influences on the surrounding tissue, potentially dictating their pathological impacts.

The mixed-type plaque—coexisting dense and loose regions—implies a dynamic interplay, supporting the hypothesis that dense and loose types represent independent growth modes that can converge (Extended Data Fig. 10b). Biologically, these modes reflect distinct pathogenic strategies: the loose-type favors diffusive, infiltrative growth, while the dense-type prioritizes volumetric expansion and area coverage. Together, these architectural strategies contribute to plaque progression through divergent mechanisms.

Our findings also identify microglia as key drivers of plaque heterogeneity. As the brain’s resident phagocytes, microglia actively interact with Aβ plaques ^24, 26, 38^, but their role in plaque biogenesis remains controversial—proposed to clear Aβ^48–50^ yet also linked to plaque compaction and propagation ^38, 51–53^. Our data resolve this paradox by revealing a direct structural impact: microglial contact drives Aβ branching. Microglial processes interdigitate with Aβ branch termini, fully encasing individual barbs. Plaque regions in contact with microglia exhibit 2.3-fold higher barb density than microglia-free regions, while “escaping” Aβ branches (not covered by microglia) are loose and unbranched. These escaping branches connect directly to Aβ coronas—structures with identical smooth morphology—suggesting coronas represent unmodified, newly formed Aβ that has not interacted with microglia.

This implies microglia tip the balance toward dense-type characteristics. While compaction may limit fibril elongation, it expands the plaque’s spatial footprint and increases local Aβ density, amplifying protofibril-rich reactive hotspots linked to neurotoxicity^54, 55^. Our data also show dense-type plaques exhibit explosive growth, in contrast to the gradual accumulation of loose-type plaques. Thus, microglia may not simply clear or compact Aβ but remodel plaque structure in ways that modulate expansion and toxicity. Resolving the molecular mechanisms of this interaction will be critical for targeting microglia in AD therapies.

More broadly, our identification of Aβ branches as the fundamental plaque unit and microglia as structural modulators reshapes our understanding of Aβ pathology. By resolving how molecular aggregates scale to tissue-level lesions, we provide a framework for testing hypotheses about Aβ spread (e.g., whether branches act as seeds for new plaques) and for developing therapies targeting plaque architecture, rather than just Aβ aggregation itself. In conclusion, PASTA enables unprecedented visualization of Aβ plaque ultrastructure, revealing the structural basis of plaque formation and microglia-mediated remodeling. These findings advance our understanding of AD pathogenesis and provide a versatile framework for future AD research.

## Acknowledgments

Human tissue was provided by the National Health and Disease Human Brain Resource Center, Medical College of Zhejiang University, Hangzhou, China. This study was supported by the China Human Brain Banking Consortium, by grants from ST12030-Major Project #20212D0201100, Task2#20212D0201102, Task6#20212D0201106. We thank Prof. Xu Li for the 3×Tg mice sample collection. We thank the Instrumentation and Service Center for Molecular Sciences, Instrumentation and Service Center for Physical Sciences, and Advanced Biomedical Technology Core Facility at Westlake University for technical support. We thank the Microscopy Core Facility of Westlake University for the facility support and thank technicians Y. Gao, F. Xiao, L. Xu, M. Liao, and F. Zhao for technical assistance. We thank F. Xu at the Westlake University High Throughput Core Facility for assistance in SPR data collection. We thank the Laboratory Animal Resources Center (LARC) at Westlake University for animal maintenance.

## Author contributions

F.W. synthesized, screened, and characterized the probes, performed FLIM and SIM imaging, processed and analyzed all imaging data, and wrote the manuscript; F.W. and X.S. performed ExM imaging; X.S. performed CLEM imaging; F.W. and Y.H. performed confocal imaging; Q.Z. and B.C. prepared cryo-EM samples, performed cryo-EM data collection, cryo-EM data processing, and atomic model building. C.L. and D.L. supervised the cryo-EM-related experiments and data processing. Y.H., L.X., and S.Z. maintained the animals; L.X. contributed to troubleshooting; D.Y. and K.D.P. supervised the project.

