## Supplementary Material for "Probe-assisted Structural Topography Reveals In-tissue Lesion Heterogeneity in Alzheimer’s Disease"

[1] School of Life Sciences, Westlake University, Hangzhou 310024, China
[2] Bio-X Institutes, Key Laboratory for the Genetics of Developmental and Neuropsychiatric Disorders (Ministry of Education), Shanghai Jiao Tong University, Shanghai, China
[3] Zhangjiang Institute for Advanced Study, Shanghai Jiao Tong University, Shanghai, China
[4] National Health and Disease Human Brain Tissue Resource Center, School of Medicine, Zhejiang University, Hangzhou, 310012, China
[5] Department of Pathology, The First Affiliated Hospital, Zhejiang University School of Medicine, Hangzhou, 310058, China
[6] Interdisciplinary Research Center on Biology and Chemistry, Shanghai Institute of Organic Chemistry, Chinese Academy of Sciences, Shanghai, China
[7] State Key Laboratory of Chemical Biology, Shanghai Institute of Organic Chemistry, University of Chinese Academy of Sciences, Chinese Academy of Sciences, Shanghai, China
[8] Shanghai Academy of Natural Sciences (SANS), Fudan University, Shanghai, China
[9] Westlake Laboratory of Life Sciences and Biomedicine, Hangzhou 310024, China
[10] These authors contributed equally

### Methods

#### Antibodies and reagents

The following antibodies were used in this study: AF488 anti-β-Amyloid 17-24 (BioLegend, 4G8, 800713, 1:100), anti-β-Amyloid (Cell Signaling, D54D2, 8243S, 1:800), anti-β-Amyloid 1-42 (Cell Signaling, E6D5M, 24090S, 1:400), anti-phospho-Tau (Ser202, Thr205) (Thermo Fisher Scientific, AT8, MN1020, 1:1000) and anti-Iba1 (Wako, 019-19741, 1:500). All secondary antibodies used were conjugated with Alexa dyes from Thermo Fisher Scientific (1:200). Fluorescently labeled $A\beta_{1-42}$ peptides were provided by Qyaobio (China). Lyophilized peptides (purity >95%) were dissolved in DMSO at a concentration of 1 mg/mL and stored as stock. Small molecule dyes used in the study are: Hoechst33342 (Thermo Fisher Scientific, H1399), Thioflavin T (Sigma Aldrich, T3516), Thioflavin T (Abcam, ab120751), Thioflavin S (Sigma Aldrich, T1892), Congo Red (Sigma Aldrich, C6277), Methoxy-X04 (MedChem Express, HY-103240), and X-34 (Sigma Aldrich, SML1954). Phosphate-buffered saline (1×PBS) mentioned in the following sections contains 2.0 mM $KH_2PO_4$, 137 mM NaCl, 10 mM $Na_2HPO_4$, and 2.7 mM KCl at pH 7.4.

#### Generation of Aβ intermediates and fibrils

$A\beta_{1-42}$ peptide was purchased from QYAOBIO. Dried peptide film was prepared following a previously established protocol [56]. In brief, the $A\beta_{1-42}$ peptide was solubilized in hexafluoroisopropanol (HFIP) at a concentration of 1 mM and incubated for 60 min at room temperature. The tube of the Aβ solution was left with the lid open overnight at room temperature to allow HFIP evaporation. The remaining traces of HFIP were evaporated on a rotary vacuum concentrator (Christ, RVC 2-18 CDplus), and the peptide was dried for 10 min. $A\beta_{1-42}$ oligomer was prepared according to Kayed's protocol [57]. Briefly, dried peptide film was re-dissolved in HFIP to a concentration of 2.5 mg/mL and incubated for 20 min at room temperature. The resulting seedless solution was diluted 10× into deionized distilled water ($ddH_2O$). After incubation for 20 min at room temperature, the sample was centrifuged for 15 min at 14,000 g, and the supernatant fraction was transferred to a new siliconized tube and subjected to a gentle stream of $N_2$ for 10 min to evaporate HFIP. The sample was agitated at 500 rpm for 24–48 h at 22°C. $A\beta_{1-42}$ protofibril was prepared according to Lashuel's protocol [58]. Briefly, dried peptide film was dissolved in DMSO (final concentration 20 mg/mL), and $ddH_2O$ was carefully added (final concentration 1.25 mg/mL). After the addition of 2 M Tris base solution (pH 7.5, final Tris concentration 25 mM), the sample was vortexed for 2–5 s and then incubated for 5 min. Mature fibrils of $A\beta_{1-42}$ were prepared by dissolving dried peptide film in DMSO to a concentration of 5 mM and immediately diluted in 1×PBS to a final concentration of 50 µM. The sample was incubated at 37 °C under gentle agitation for 7 days. The formation of the above Aβ species was confirmed by AFM.

#### Absorbance and fluorescence measurements

Absorbance and fluorescence spectra of the testing solutions were recorded with a UV-visible spectrometer (Agilent Cary3500) and a spectrofluorometer (Edinburgh FS5), respectively. Probes were dissolved in DMSO to make a 20 mM stock solution, which can be diluted to the required concentration of testing solution for measurement.

Aliquots of protein solutions were added to the probe testing solutions to their desired concentrations with gentle mixing. Absorbance and fluorescence of the testing solutions were recorded after 5 min of incubation.

**Viscosity sensitivity measurements**

Viscosity sensitivity was measured according to published protocols [21, 59]. Probes are diluted from a 20 mM DMSO stock into a series of ethylene glycol and glycerol mixtures at 20 °C to a final concentration of 10 μM. The composition and viscosity of each mixture at 20 °C are described in the table below. The emission spectra of sample solutions were collected with a fluorescence spectrophotometer (Hitachi, F-7100), and the emission peak intensity was plotted as a function of solvent viscosity in a double logarithmic manner and fitted by linear regression. The slope of the regression line represents the viscosity sensitivity value.

| Composition Glycerol vol% | 30% Glycerol | 50% Glycerol | 60% Glycerol | 70% Glycerol | 38% Glycerol |
|---|---|---|---|---|---|
| Viscosity (mPa·s) | 81 | 183 | 283 | 426 | 621 |

**Time-resolved photoluminescence (TRPL) spectra measurements**

Time-resolved photoluminescence (TRPL) spectra measurements were made on an Edinburgh Instruments, FLS1000 Photoluminescence Spectrometer, with a 635 nm EPL laser source (less than 100 ps, 2 MHz). Fluorescence decay traces were analyzed using Fluoracle software. The decays were fitted to biexponential functions for better $\chi^2$ values. The quality of the fits was assessed based on the $\chi^2$ values and residuals distribution.

**Aggregation assays**

Monitoring of Aβ aggregation was performed adapted from previously established protocols [60, 61]. The aggregation solution containing 100 μM $A\beta_{1\text{-}42}$ in 1×PBS (pH 7.4) was shaken at 600 rpm at 37 °C. At indicated time points, fluorescence spectra were obtained by mixing a 10 μL aliquot of aggregation solution with probe (NapIM or ThT) in 1×PBS to make a final concentration of 10 μM probes with 20 μM Aβ. At certain time points, a 10 μL aliquot of aggregation solution was spotted onto freshly cleaved mica, washed, and dried for later AFM acquisition. ThT signal was monitored at 440 nm excitation and NapIM signal at 607 nm excitation. Each sample was run in triplicate.

**Atomic force microscopy**

AFM experiment was carried out following a previously established protocol [62, 63]. Samples containing Aβ were prepared by spotting 10 μL of Aβ solution onto freshly cleaved mica (Ted Pella, Inc.). Droplets were incubated on the substrate for 10 min at room temperature, washed twice with $ddH_2O$, and dried under a $N_2$ gas stream. Images were obtained using a Jupiter XR Atomic Force Microscope using a Z scanner with a cantilever holder with an xy range of 100 μm. Tapping mode was employed for all images using a silicon cantilever with a silicon tip (AC160TS-R3, Olympus) with a tip radius of 9 ± 2 nm. At least four regions of the mica surface were examined to ensure that similar structures existed throughout the sample. The images were processed using Gwyddion (version 2.62).

**Transmission electron microscopy (TEM)**

5 μL of sample solution was placed onto carbon coated grids (BZ11033a, EMCN). The sample was incubated on the grid for 10 min at room temperature. Excess solvent was soaked away with filter paper. The grid was washed once with 10 μL water and stained once with 10 μL 1% w/v phosphotungstic acid (PTA) in water for 1 min. Excess PTA solution was soaked away with filter paper and the grid was dried overnight. The dried grid was examined in 120 kV TEM (Talos L120C G2).

**Co-precipitation of NapIM with Aβ fibrils**

Aβ fibrils (50 μM) were incubated with NapIM (50 μM) in 0.5 mL test tubes at 25 °C for 1 h in the dark. Tubes with 50 μM probe alone and fibrils alone were set as controls. After centrifugation at 21,000 g for 30 min, coprecipitation of NapIM with Aβ fibrils was observed owing to the color of NapIM. Fluorescent imaging of test tubes was acquired on the ChemiDoc MP Imaging System (BioRad). Images of tubes without centrifugation were also recorded under the same acquisition run. The experiment was carried out in the dark.

**Fluorescent binding assays**

Solutions with various concentrations of probes were obtained by adding different amounts of probes to the PBS solution of Aβ (5 μM). Fluorescence intensity was then determined in a Thermo plate reader using 607/711 (NapIM) or 440/480 (ThT) excitation/emission filters. The saturation binding curve and $K_d$ value were analyzed by GraphPad Prism 10 with non-linear one-site binding regression.

**SPR binding assays**

Protein solutions were diluted into sodium acetate buffer (pH 4.0) and immobilized on a CM5 sensor chip (Cytiva). A serially diluted probe (NapIM or ThT) solution was flowed over the chip in HEPES buffer (10 mM HEPES, 150 mM NaCl, pH 7.4, 0.05%Tween-20). Binding affinities were measured using a Bioacore 8K+ (Cytiva) at 25 °C in the multi-cycle mode. Binding kinetics were analyzed with Biacore evaluation software (Cytiva) to calculate the $k_a$ and $k_d$ values. The $K_D$ values were calculated as the ratio of ligand dissociation rate to association rate ($k_d$/$k_a$).

**Postmortem human brain sample**

Postmortem brain tissue was obtained from three clinically diagnosed Alzheimer's disease (AD) cases. Ethical approval for this study was granted by the Institutional Review Board of the National Health and Disease Human Brain Resource Center, Medical College of Zhejiang University (2020-005). Human tissue was provided by the National Health and Disease Human Brain Resource Center, Medical College of Zhejiang University, Hangzhou, China.

Among them, frontal lobe tissue from a 78-year-old female (case ID: 2023CBB079) was used for pathological and molecular analyses. Neuropathological examination revealed AD-related changes classified as A3B3C3 (Thal phase 5, Braak stage VI), with coexisting TDP-43 proteinopathy (LATE stage), α-synuclein pathology (Braak stage IV), cerebral amyloid angiopathy (CAA stage 1), and basal ganglia arteriosclerosis. Occipital lobe tissue from a 78-year-old male (case ID: 2020021) was used for immunofluorescence and co-staining with NapIM. Neuropathological examination revealed AD-related changes classified as A2B1C1 (Braak stage 1) with coexisting CAA, and the genotype for APOE was e3/e4. Clinical diagnoses included aspiration pneumonia, Alzheimer's disease, primary hypertension, chronic enteritis, bradyarrhythmia, cholelithiasis with cholecystitis, and anemia. All tissue collection and processing followed standardized protocols and ethical guidelines.

**Amyloid fibril extraction**

Amyloid-β (Aβ) fibril was extracted from postmortem frozen brain tissue of an individual with AD following previously reported protocols with moderate modification. Briefly, frozen frontal lobe tissue was manually homogenized in 20 volumes (v/w) of extraction buffer containing 10 mM Tris-HCl (pH 7.5), 0.8 M NaCl, 10% sucrose, 1 mM EGTA, 0.1% sarkosyl, 0.2 M PMSF, and protease inhibitor cocktail (Roche). The homogenate was adjusted to a final concentration of 2% sarkosyl and incubated at 37 °C for 1 h with constant mixing on a rotary mixer. Following incubation, the homogenate was centrifuged at 10,000 *g* for 10 min at 4 °C. The supernatant was subjected to ultracentrifugation at 100,000 *g* for 60 min at 4 °C using SW41Ti (Beckman Coulter). The supernatant fraction was then further centrifuged at 166,000g for 1 h at 4 °C using the same rotor. The final pellet was resuspended with 20 mM Tris-HCl, pH 7.4 (100 µl/g tissue).

**Immunogold NS-TEM**

The brain-extracted fibrils were diluted 4-fold and applied to a glow-discharged carbon-coated copper grid (200 mesh), followed by incubation for 10 min. The grid was then blocked with 0.1% BSA for 30 min at room temperature, followed by incubation with primary antibody against Aβ (NAB228, Cell Signaling Technology; 1:10 dilution) for 30 min at room temperature. After three washes with PBS, the grid was incubated with 6 nm colloidal gold conjugated anti-mouse secondary antibody (1:15; 711-205-152, Jackson ImmunoResearch) for 30 min at room temperature. The grid was sequentially washed with PBS, double-distilled $H_2O$, and 2% w/v uranyl acetate, and then stained with 2% w/v uranyl acetate for 45 s. Excess buffer was removed with filter paper. Transmission electron microscopy (TEM) micrographs were acquired using a Tecnai T12 microscope (FEI) operated at 120 kV.

**Cryo-EM sample preparation and data collection**

To prepare fibril samples with chemical ligand or DMSO, the fibrils were diluted fourfold and incubated with 150 µM NapIM or 1.5% DMSO in the presence of pronase for 1 h. The 4-µl aliquot of fibril-containing solution was applied to a glow-discharged holey carbon-coated grid (C-Flat, 300-mesh, 1.2/1.3; 71159), incubated for 8 s at 16 °C and 95% humidity, and then blotted with filter paper. The grid was subsequently plunge-frozen in liquid ethane using Vitrobot Mark IV (FEI, Thermo Fisher Scientific).

For image acquisition, the sample was imaged on a Thermo Fisher Titan Krios G4 cryo-TEM instrument (Thermo Fisher Scientific) at 300 kV, equipped with a BioContinuum K3 direct detector (Gatan) in counting mode. A GIF Quantum energy filter (Gatan) with a 20 eV slit width was used to remove inelastically scattered electrons. The super-resolution videos were recorded at ×105,000 magnification, yielding a pixel size of 0.83 Å, with a total dose of ~55 $e^-/Å^2$ over a 2.0 s exposure. Automated cryo-EM data collection was performed using EPU software (Thermo Fisher Scientific), with defocus values ranging from -1.0 to -2.4 µm.

**Cryo-EM image pre-processing and helical reconstruction**

For image pre-processing, 40 movie frames per micrograph were corrected for beam-induced motion, aligned, dose-weighted, and further binned with a physical pixel size of 0.83 Å using MotionCorr2. The resulting micrographs were then used for contrast transfer function estimation with CTFFIND-4.1.8. Helical reconstruction was subsequently carried out in RELION 3.1.

**Apo Aβ fibrils dataset**

Fibrils were manually picked using the "Manual picking" program in RELION 3.1 and initially extracted to segments with a box size of 288 pixels, with an inter-box distance of 23.9 Å. These particles were subsequently re-extracted using a larger box size of 1050 pixels and downscaled to 512 pixels. The reference-free 2D classification steps were then performed with a decreasing in-plane angular sampling rate from 2 to 0.5 and a T = 2 regularization parameter to calculate the apparent half pitches from the 2D class averages and to discard the segments contributing to suboptimal 2D class averages. For type I $A\beta_{42}$ fibrils, an initial 3D model was constructed de novo from purified segments spanning an entire helical crossover, using the relion_helix_inimodel2d program. The purified segments were re-extracted with 288 pixels. These segments and the initial 3D model that was low-pass-filtered to 30 Å were further applied to perform 3D classifications (k = 3) using the helical parameters calculated through the splicing of 2D class averages. The best 3D reconstruction map was selected and subjected to an additional round of 3D classifications (k = 1) with local optimization of helical twist and rise, during which β-strands perpendicular to the helical axis became clearly resolved. Then, 3D auto-refinement with further optimization of helical parameters was performed. To enhance resolution, Bayesian polishing and CTF refinement were carried out, followed by additional rounds of 3D autorefinement. Finally, the map was sharpened with a soft-edge solvent mask using the standard "post-processing" program in RELION 3.1. Overall resolution was estimated based on the gold-standard 0.143 Fourier shell correlation (FSC) between the two independently refined half-maps. Then the local resolution was estimated using the "Local resolution" procedure in RELION 3.1, applying the same mask and B-factor in post-processing.

**NapIM-Aβ complex dataset**

Fibrils were manually picked using the "Manual picking" program in RELION 3.1 and initially extracted to segments with a box size of 288 pixels, with an inter-box distance of 23.9 Å. These particles were subsequently re-extracted using a larger box size of 960 pixels and downscaled to 512 pixels. The reference-free 2D classification steps were performed with a decreasing in-plane angular sampling rate (from 2 to 0.5) and a T = 2 regularization parameter to determine the apparent half pitches from the 2D class averages and to discard the segments contributing to suboptimal 2D class averages. For type I $A\beta_{42}$ fibrils, an initial 3D model was generated de novo from purified segments spanning an entire helical crossover, using the relion_helix_inimodel2d program. The purified segments were re-extracted with 288 pixels and used for 3D classifications (k = 3), employing helical parameters derived from 2D class average splicing. The best 3D reconstruction map was selected and subjected to 3D autorefinement with further optimization of helical parameters. To further improve resolution, Bayesian polishing and CTF refinement were performed, followed by another round of 3D autorefinement. The final density map was sharpened using a soft-edge solvent mask with the standard “post-processing” program in RELION 3.1. Overall resolution was estimated based on the gold-standard 0.143 Fourier shell correlation (FSC) between the two independently refined half-maps. Local resolution was then estimated using the “Local resolution” procedure in RELION 3.1, applying the same mask and B-factor settings used during post-processing.

**Structural model building and refinement**

The starting model of type I $A\beta_{42}$ fibrils from AD was based on the previously determined structure of the type I $A\beta_{42}$ fibril (Protein Data Bank (PDB) 7Q4B). The starting three-layer coordinate model of the fibril core was manually fitted into the central region of the sharpened density map using UCSF Chimera 1.13.1, followed by manual adjustment in WinCoot 0.8.9. Real-space refinement was performed using the phenix.real_space_refine program in PHENIX 1.13, with secondary structure and geometry restraints applied. The final model was validated by the ‘comprehensive validation’ procedure in PHENIX 1.13.

For the NapIM-Aβ complex, the coordinates and geometry restraints of NapIM were generated using the SMILES string in phenix.elbow. The ligand was manually docked into the central region of the sharpened density map using Chimera 1.13.1. Simultaneously, the $A\beta_{42}$ model was manually adjusted in WinCoot 0.8.9.2 and refined in PHENIX 1.13 using phenix.real_space_refine program, applying rotamer, Ramachandran, and geometry restraints. Additional details for helical reconstruction and model building were summarized in Supplementary Table 1.

**Animals and animal care**

All mouse maintenance and experimental procedures complied with the Institutional Animal Care and Use Committee guidelines of the School of Life Sciences, Westlake University (approval no. AP#22-053-2-YD-14). 5×FAD (Stock No. 034848-JAX) were kindly provided by the laboratory of Danyang He (Westlake University) and bred in the Laboratory Animal Resources Center of Westlake University. Cx3cr1-2A-EGFP mice were purchased from Shanghai Model Organisms (NM-KI-210099). Cx3cr1-2A-EGFP/5×FAD +/+ mice were produced by crossbreeding 5×FAD with Cx3cr1-2A-EGFP mice. All mice were bred in specific-pathogen-free conditions, maintained on a 12-h light/dark cycle at 22°C with 40%–60% humidity. Standard chow and water were provided to the mice ad libitum. Both male and female mice were used in the study, ranging in age from 2 months to 13 months old. The specific age of the mice used in each experiment was described in the text and corresponding figure legends.

**Fixed brain sample preparation**

Mice were euthanized and perfused with 1×PBS followed by 4% paraformaldehyde (PFA). Brains were dissected and fixed in 4% PFA overnight at 4 °C. These brains were then sectioned at 40 µm in thickness using a Leica vibratome (VT1200S) and stored at 4 °C in PBS + 0.05% ProClin 300.

**Fresh brain slice Aβ labeling**

Fresh brain labeling was carried out following a previously established protocol [24].

Mice were decapitated under anesthesia. The brains were dissociated with cooled ACSF with continuous oxygenation. After dissociation, the brains were immediately transferred onto a vibratome. Coronal sections were made at 150-µm thickness. Fluorescently labeled A$\beta_{1-42}$ (FAM-A$\beta_{1-42}$, QYAOBIO) was added to the incubation bath to make a final concentration of 2 µg/mL. Slices were incubated on a shaker for 1 h at 4 °C and then washed twice with ACSF. After the last wash, slices were fixed in 4% PFA at room temperature for 1 h. Fixed slices were then washed with PBS before imaging.

**Immunohistochemistry**

The fixed brain slices were permeabilized with 0.5% Triton-X 100 in PBS for 30 min, and washed with 1×PBS for 3 times. These brain slices were pretreated with 70% formic acid in 1×PBS for 10 min and washed with 1×PBS for 5 min. Samples were blocked with 4% (v/v) goat serum (Gibco, 16210064) in 1×PBS with 0.2% (v/v) Triton-X-100 for 1 h at room temperature. Primary antibodies diluted in blocking solution were applied to the sample slices at 4 °C overnight. Brain slices were washed 3 times in 1×PBS and then incubated in matched secondary antibodies at room temperature for 1 h. Samples were washed 3 times in 1×PBS. For ThS staining, samples were incubated with ThS (0.273 g/L) in 50% ethanol for 8 min, followed by two 3-min washes with 50% ethanol at room temperature. For NapIM and P2–P5 staining, samples were incubated with a probe (20 µM) in 1×PBS for 1 h. All samples were incubated with Hoechst33342 (5 µg/mL) in 1×PBS for 1 h to label nuclei. Stained sections were mounted on glass slides with mounting medium (SouthernBiotech, Fluoromount-G®, 0100-01). Mounted slides were imaged immediately or stored at 4 °C until imaged.

**Fluorescence lifetime imaging microscopy (FLIM)**

FLIM experiments were performed using a Leica STELLARIS 8 FALCON confocal microscope, equipped with an adjustable pulsed white light laser and a 100× oil immersion objective lens (Leica, Microsystems). The brain sections stained with NapIM were mounted on the glass slide. The laser was tuned to 653 nm for NapIM excitation. Emission was detected within the wavelength ranges of 666–829 nm. Images were captured in a 1024×1024 format with three frames average. Z-stack NapIM multiplexing was performed with the 17 µm z-scan at a 500 nm step size. NapIM staining patterns were separated through the FLIM mode analysis or phasor plot analysis using the LAS X FLIM software (Leica Microsystem). The phasor plot analysis module transforms the recorded fluorescence lifetime decay data [64]. The average lifetime values were exported from the LAS X FLIM software (Leica Microsystems).

**3D-SIM imaging of Aβ plaques**

3D-SIM images of NapIM-labeled Aβ plaques were acquired on the Multi-SIM imaging system (NanoInsights) with a 100×/1.49 NA oil objective (Nikon CFI SR HP Apo), solid-state lasers (640 nm), and a sCMOS (Complementary Metal-Oxide-Semiconductor) camera (Photometrics Kinetix). Serial z-stack sectioning was done at 135 nm intervals for 3D-SIM mode. Immersion oils with refractive indices of 1.518 were used for brain slices mounted on glass slides. The microscope is routinely calibrated with 100 nm fluorescent spheres to calculate the lateral and axial limits of image resolution. SIM image stacks were reconstructed using SI-Recon 2.23.3 (NanoInsights) with the following settings: pixel size 30.6 nm; channel-specific optical transfer functions;

Wiener filter constant 0.005 for 3D mode; discard Negative Intensities background. Then the reconstructed SIM image was denoised with a total variation (TV) constraint.

**Correlative light and electron microscopy (CLEM)**

TEM-CLEM (Correlative light and electron microscopy) was used to validate the ultrastructure of NapIM staining patterns in brain sections. Fluorescence images were acquired before the tissue slices were prepared for TEM. Briefly, following the deep anesthesia, the female 9-month-old 5×FAD mouse was transcardially perfused with sodium cacodylate buffer (0.1 M, pH = 7.4) and cold 4% PFA (Electron Microscopy Sciences, US), and the brain tissue was harvested and post-fixed at 4 °C. Coronal brain sections with a 100 µm thick were sliced using a VT1200S Vibratome. After the routine staining, the z-stack images of NapIM and DAPI staining patterns at high resolution were acquired using the Nikon Spinning Disk microscope with the 60x oil objective lens. Following imaging, the slices were incubated with mixed fixation reagent (2.5% glutaraldehyde and 2% PFA in 0.1 M sodium cacodylate buffer) at 4 °C overnight and washed three times for 30 mins in ice-cold 0.1 M sodium cacodylate buffer. Next, the slices were postfixed in ice-cold reduced 1% osmium tetroxide (Ted Pella, USA) in cacodylate buffer for 1 h (protected from light) and rinsed with 0.1 M sodium cacodylate buffer three times. After washing with cold-dd$H_2O$ three times, the slices were stained with 1% uranyl acetate in dd$H_2O$ at room temperature (22–25°C) for 1 h in the dark. The slices were further rinsed with chilled dd$H_2O$, dehydrated with a graded ethanol series (30%, 50%, 70%, 90%, and 100% three times, respectively), and flat-embedded with Durcupan ACM resin. Finally, followed by at least 48 h polymerization at 70°C, ultrathin sections with 70 nm thickness were prepared. According to the fluorescent z-stack images, the ultra-field of plaques at the same regions was captured at different magnifications using a transmission electron microscope (Thermo Scientific, Talos L120C G2) at 80 kV.

**Expansion microscopy**

Sources and identifiers for the chemicals and reagents used for expansion microscopy are listed in Supplementary Table 2. Based on testing with several published expansion microscopy methods [65, 66], the Magnify protocol [67] was used for the ExM imaging of NapIM staining. Briefly, a monomer solution (4% (v/v) DMAA, 34% (w/v) SA, 10% (w/v) AA, 0.01% (w/v) Bis, 1% (w/v) NaCl, and 1× PBS) was prepared before use. The brain sections were flattened at the bottom of the gelation chamber and incubated with freshly prepared activated monomer solution (added with a final concentration of 0.25% (w/v) APS, 0.2% (v/v) TEMED, 0.001% (w/v) 4-HT, and 0.1% (v/v) methacrolein) at 4 °C for 30 min. A trimmed glass slide was placed to cover the gelling chamber, and the samples were then polymerized in a humidified container under nitrogen at 37 °C for 3 h. The gelled tissue sections were carefully detached from the glass slides and immersed in the digest buffer (50 mM Tris, 1 mM EDTA, 0.5% Triton X-100, 1 M NaCl, pH = 8.0) supplemented with Proteinase K with a final concentration of 2–4 U/mol for 2–4 h at 22–25°C. The homogenized samples were quickly washed three times with 1×PBS at 22–25°C and then incubated in NapIM staining solution with the final working concentration of 40 µM in 1×PBS at 4°C for 2–3h, counterstained with DAPI as well. Then the samples were washed three to four times for at least 10 min each with 1×PBS at 22–25°C and then immersed in $ddH_2O$ for 2–3 h to expand. This step was repeated 3–5 times in fresh $ddH_2O$ until the size of the expanding hydrogel sample plateaued (up to 6.5). The expanded samples were imaged and stored at 4°C if further needed. The dimensions of the same brain sections were measured before and after expansion. The calculated ratio of the expanded diameter over the original diameter was defined as the linear expansion factor for image calibration.

The brain tissue sections before expansion were imaged using a wide-field Nikon Eclipse Ti2-E inverted microscope. After expansion to the targets, the large z-stack imaging of plaques was performed using the Nikon Spinning-Disk Field Scanning Confocal System (with CSU-W1 SoRa imaging setup) with the 40× NA1.15 water objective lens. The fractal dimension ($D_f$) of the expanded plaque structures was quantified using the "Fractal Box Count" plugin in Fiji (ImageJ), applied to binary masks of the segmented Aβ signals at the maximum cross-sections of plaque 3D images. 3D reconstruction was performed using Imaris 10.1.0. Manual segmentation of plaque structures was conducted using VAST v.1.5.0 (downloaded from https://lichtman.rc.fas.harvard.edu/vast/).

**Classification of plaque subtypes**

The three stages (D1, D2, D3 for dense type and L1, L2, L3 for loose type) were identified based on morphological progression with criteria shown below:

| Type | Dense Type | | |
|---|---|---|---|
| Stages | D1 (Early) | D2 (Intermediate) | D3 (Late stage) |
| Size | minFeret < 5 µm | minFeret 5–8 µm | minFeret > 8 µm |
| Architecture | Tightly packed architecture. | Tightly packed architecture. | Tightly packed architecture. |
| Type | Loose Type | | |
| Stages | L1 (Early) | L2 (Intermediate) | L3 (Late stage) |
| Size | minFeret < 5 µm | minFeret 5–8 µm | minFeret > 8 µm |
| Architecture | Loosely arranged branches without spikey sprouts. Mostly linear morphology, might show stem-and-branch architecture. | Loosely arranged Aβ branches without spikey sprouts. Showing a prominent central stem and a complete absence of compact regions. | Loosely arranged branches without spikey sprouts. |

**Statistical analysis**

Statistical analyses were carried out using Prism (GraphPad). Two-tailed Student's *t*-tests were used to analyze significant differences between the two groups, and one-way ANOVA was used for multiple comparisons. *P* values below 0.05 were considered significant, *ns*: not significant, $^{*}P < 0.05$, $^{**}P < 0.01$, $^{***}P < 0.001$, $^{****}P < 0.0001$.

### Extended Data Figures

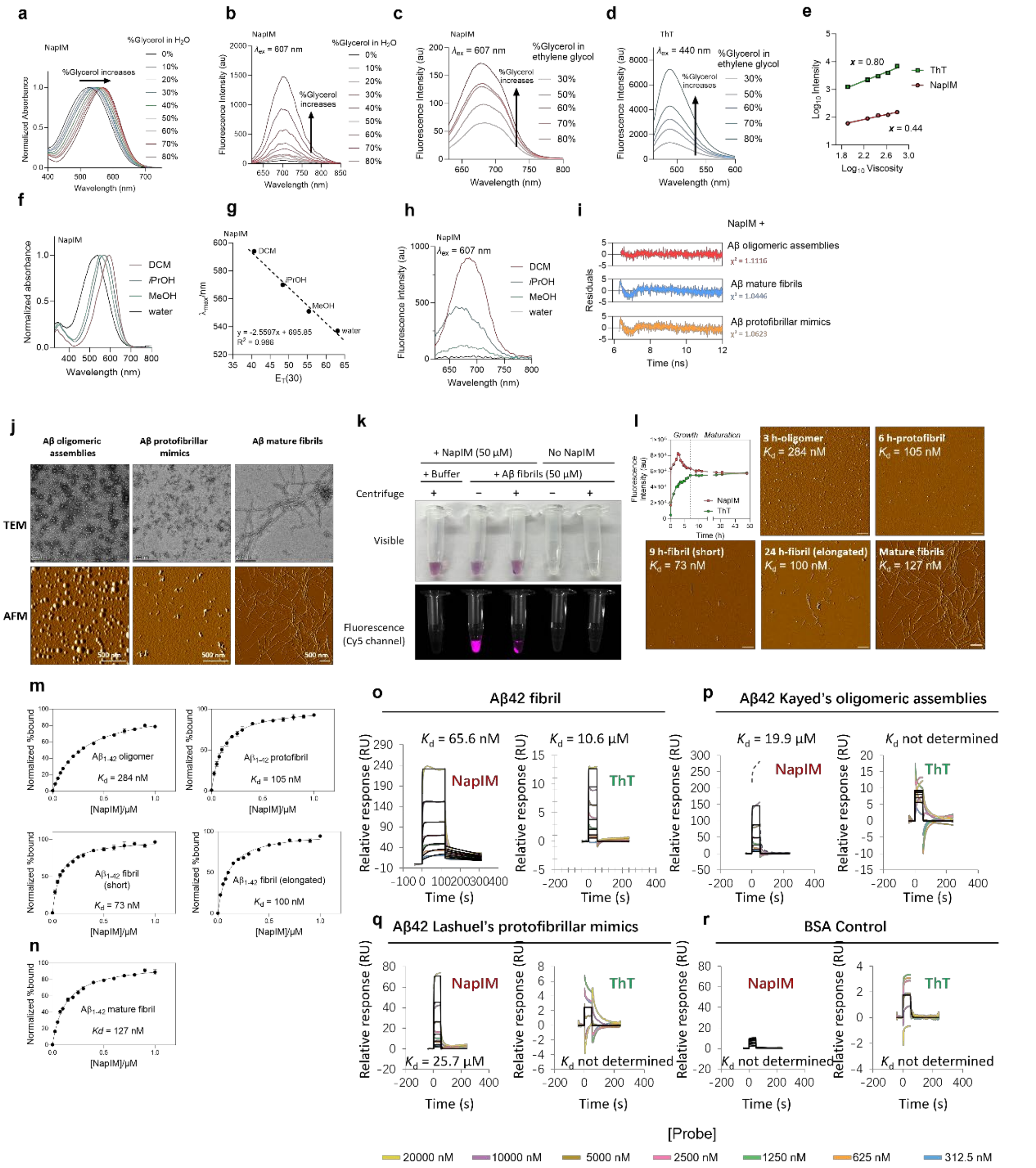

**Extended Data Fig. 1 | Characterization of NapIM.** **a**, Normalized absorbance spectra of NapIM (10 µM) in glycerol/water mixture with increasing glycerol fraction (% glycerol) at 25 °C. **b**, Fluorescence emission spectra of NapIM (10 µM) in glycerol/water mixture at increasing glycerol fraction (% glycerol) at 25 °C. **c**–**e**, Fluorescence emission spectra of NapIM (10 µM) (**c**) and ThT (10 µM) (**d**) in a series of glycerol/ethylene glycol mixture and viscosity sensitivity (**e**) calculated based on the defined viscosities [21]. **f**, Normalized absorbance spectra of NapIM (10 µM) in solvents with different polarity. **g**, Relationship of maximum absorption wavelengths against solvent polarity $E_T(30)$ Reichardt parameters [68]. **h**, Fluorescence emission spectra of NapIM (10 µM) in solvents with different polarity. **i**, The residuals of the fluorescence decay models in Fig. 1d. **j**, TEM and AFM images of Aβ42 oligomeric assemblies prepared from Kayed's method (left), Aβ protofibrillar mimics prepared from Lashuel's method (middle), and mature fibrils (right). All scale bars, 500 nm. **k**, The co-precipitation of NapIM with Aβ fibrils was observed under visible and fluorescence after centrifuge. NapIM (50 μM) was incubated with Aβ fibrils (50 μM) for 1 h at room temperature before centrifugation. **l**, Left, aggregation kinetics of Aβ42 as measured by NapIM and ThT (10 μM probes mixed with 20 μM Aβ; ThT Ex/Em 440/485, NapIM Ex/Em 607/722). Right and bottom, AFM images of Aβ42 at different time points. Dissociation constants ($K_d$) are indicated. Scale bars, 500 nm. **m**, Binding curves of NapIM to different on-pathway stages of Aβ42 in aggregation determined by fluorescent titration. Dissociation constants ($K_d$) are indicated. Error bars represent the s.e.m. calculated from three replicates (three distinct samples). **n**, Binding curves of NapIM to Aβ42 mature fibril determined by fluorescent titration. Dissociation constants ($K_d$) are indicated. Error bars represent the s.e.m. calculated from three replicates. **o**–**r**, SPR sensorgrams of a titration series of NapIM and ThT to immobilized Aβ of different forms showing the real-time interaction between probes and Aβ. Capture of NapIM by Aβ species and no binding of ThT on Aβ were observed.

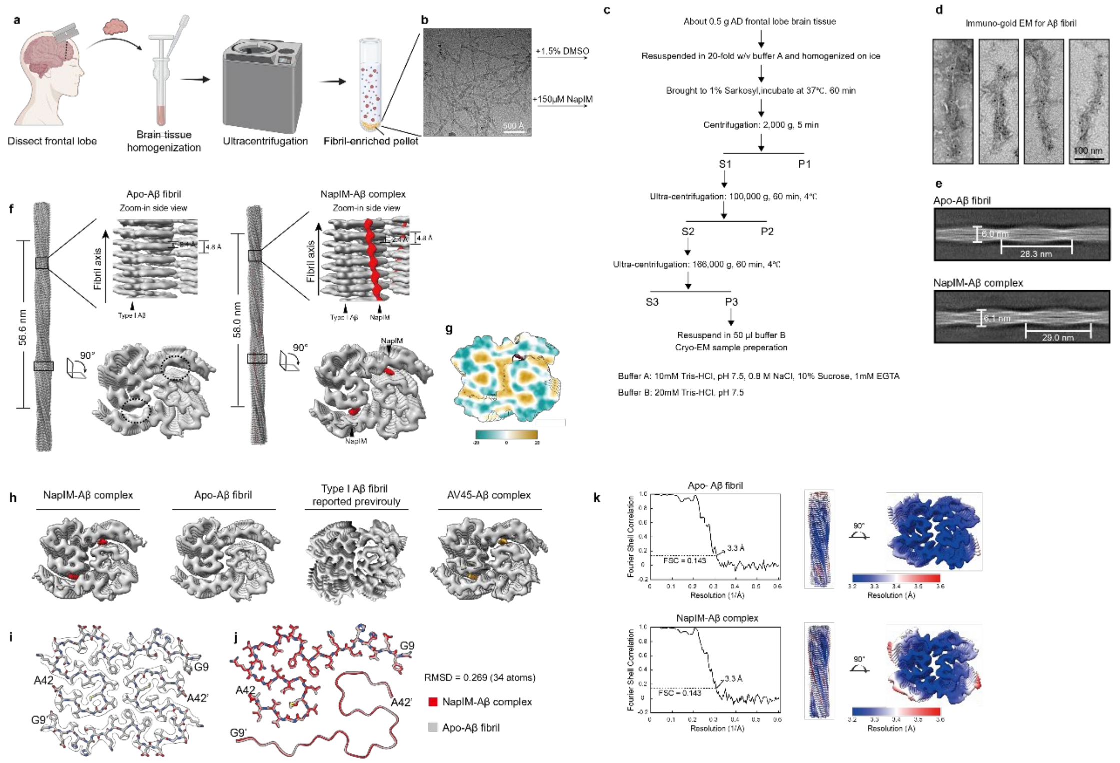

**Extended Data Fig. 2 | Structural characterization of *ex vivo* Aβ$_{42}$ fibrils after incubating with NapIM. a**, Schematic illustration of the Aβ fibril extraction method. Created with BioRender.com. **b**. Cryo-EM micrograph of the extracted fibrils, which were subsequently incubated with the NapIM molecule and the same equivalent of DMSO. **c,** Workflow of the sarkosyl-based amyloid fibril extraction procedure. Solid arrows indicate the steps used in the next steps. Fraction P3 was used for cryo-EM analysis. **d,** Representative immunogold-labeled negative-stain electron microscopy images of Aβ fibrils extracted from the frontal lobe of the AD patient. Fibrils were labelled with the antibody NAB228. Scale bar, 100 nm. **e,** The 2D class averages of fibrils in apo-Aβ fibrils and NapIM-Aβ complex. The fibril's width and crossover distance (pitch) were indicated. **f**, Cryo-EM density map of the NapIM (right panel) and apo-Aβ fibril (left panel). The maps of one crossover (360° helical turn) are shown, with zoomed-in side views and cross-section views. The helical parameters are indicated. Extra densities are colored in red for the NapIM-Aβ complex, where the same locations for the apo-Aβ fibril are circled in a dashed line. **g**, The hydrophobic surface of the Aβ fibril backbone is shown. **h,** Cross-sectional views of cryo-EM density maps from NapIM-Aβ complex, apo-Aβ fibrils, previously reported *ex vivo* $A\beta_{42}$ fibrils from sporadic AD brain, and AV45-Aβ complex. Aβ densities are shown in gray; ligand densities for NapIM and AV45 are shown in red and orange, respectively. **i,** Structural model of apo-Aβ fibrils fitted into its corresponding cryo-EM density map. Then, the density is restricted to a 2-Å radius around the structural model. **j,** Structure comparison of the type I $A\beta_{42}$ fibrils from the apo-Aβ fibrils (gray) and NapIM-Aβ complex (purple). The structural models are shown in stick and cartoon representations. Global alignment of residues 9-42 yields an r.m.s.d of 1.674 Å over 68 Cα atoms. **k**, Fourier shell correlation (FSC) curves (left) represent the gold-standard Fourier shell correlation between two independently refined half-maps, with the 0.143 criterion used to estimate the overall resolution. The corresponding local resolution maps (right) were calculated using the "Local resolution" tool in RELION 3.1.

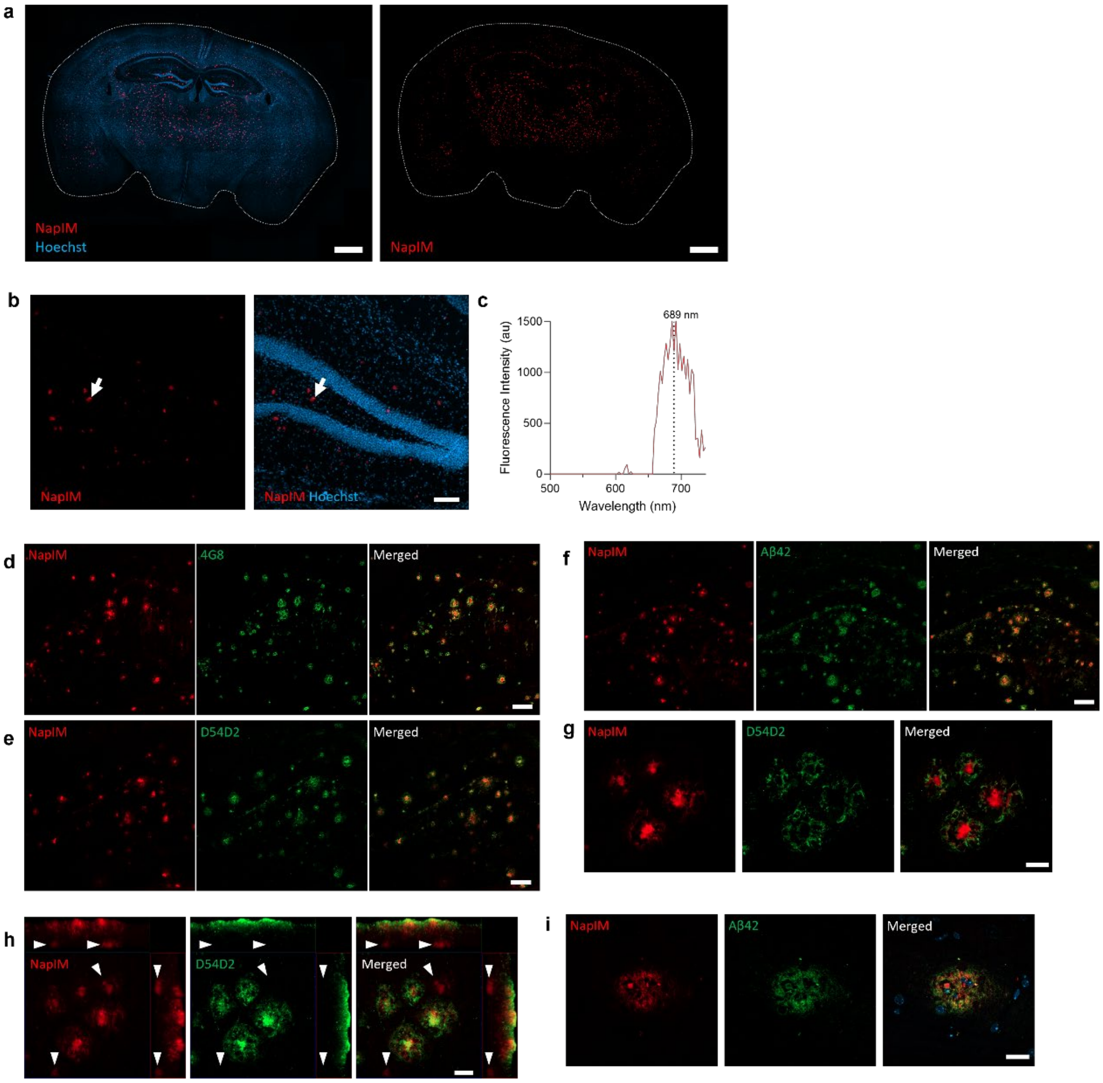

**Extended Data Fig. 3 | Staining of brain slice with NapIM**. **a**, Representative image of a 10-month-old 5×FAD mouse with NapIM and Hoechst. Scale bar, 1 mm. **b**, Images from a 10-month-old 5×FAD mouse. The white arrow indicates the plaque selected for the acquisition of the emission spectrum. Scale bar, 100 μm. **c**, Emission spectrum of NapIM-stained Aβ plaque picked from (**b**). **d**–**f**, Representative confocal images of NapIM-labeled Aβ plaques, Aβ/APP (4G8) (**d**), pan-isoform Aβ (D54D2) (**e**), and Aβ42 (**f**) from a 13-month-old 5×FAD mouse. Scale bars, 100 μm. **g**, Representative magnified images of NapIM-labeled Aβ plaques and pan-isoform Aβ (D54D2) from a 13-month-old 5×FAD mouse. Scale bar, 20 μm. **h**, MIP projections of confocal z-stack images of Aβ labeled by NapIM and pan-isoform Aβ antibody (D54D2), the same region as in **g**. Aβ plaques buried deep in tissue, only labeled by NapIM, were indicated by white arrows. Scale bar, 20 μm. **i**, Representative magnified images of Aβ plaques labeled by NapIM and Aβ42-specific antibody from a 13-month-old 5×FAD mouse. Scale bar, 20 μm.

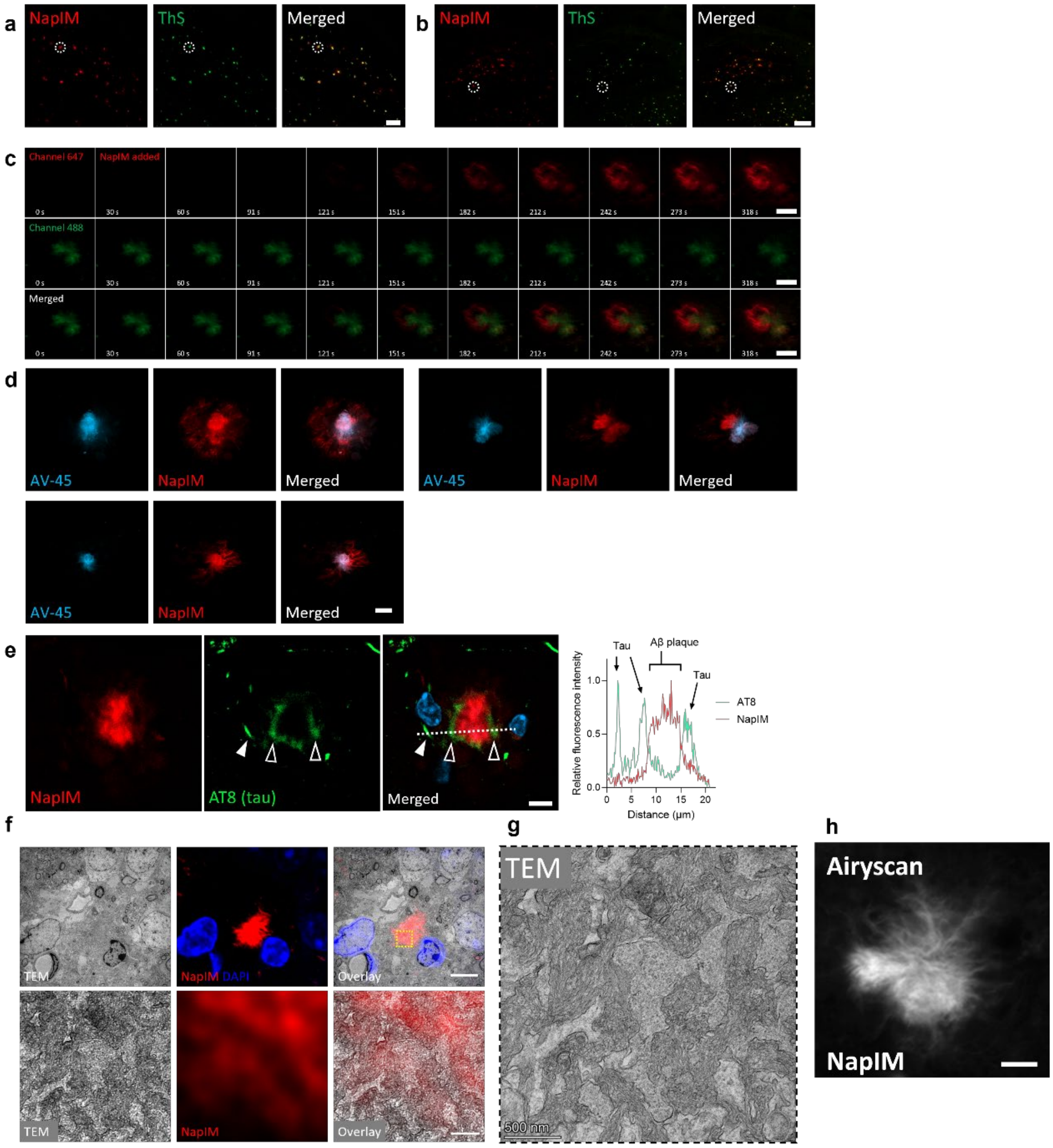

**Extended Data Fig. 4 | Validation of staining target of NapIM. a–b**, Representative confocal images of Aβ plaques labeled by NapIM and ThS from 9-month-old (**a**) or 13-month-old (**b**) 5×FAD mice. Dashed circles in (**a**) and (**b**) indicate the plaques in Fig. 2c and Fig. 2d, respectively. Scale bars, 100 µm (**a**), 200 µm (**b**). **c**, Time-lapse monitoring of an Aβ plaque pre-stained with ThS and incubated with NapIM staining solution (20 µM in PBS) from 10-month-old 5×FAD mice. Scale bars, 10 µm. **d**, Representative fluorescence images of an Aβ plaque labeled by NapIM and AV-45 from a 12-month-old 5×FAD mouse. Scale bar, 10 µm. **e**, Representative single-layer confocal images of NapIM and phospho-Tau (labeled by AT8) from a 21-month-old 3×Tg mouse. Right, line profiles of fluorescence intensities across the plaques that are shown in **e**, showing NapIM is not colocalized with tau threads (closed arrowhead) or tau material around plaque (open arrowhead). **f**, Top, representative correlative light and electron microscopy (CLEM) images of Aβ plaques from a 9-month-old 5×FAD mouse showing NapIM staining patterns colocalized with amyloid protein compositions identified by TEM. Bottom, magnified view of NapIM-labeled Aβ plaque (yellow box in top panel) with corresponding region in TEM. The NapIM signal was adjusted to a lower threshold than the top panel to show fine structure. Scale bars, 5 µm (top); 500 nm (bottom). **g**, Highly resolved TEM of the region of the yellow box in the top panel showing fibril bundles of amyloid. Scale bar, 500 nm. **h**, Representative confocal image (Airyscan mode, Zeiss LSM 980) of an Aβ plaque labeled by NapIM showing fiber-like structure from a 10-month-old 5×FAD mouse. Imaging objective, 63× Oil NA1.4. Scale bar, 2 µm.

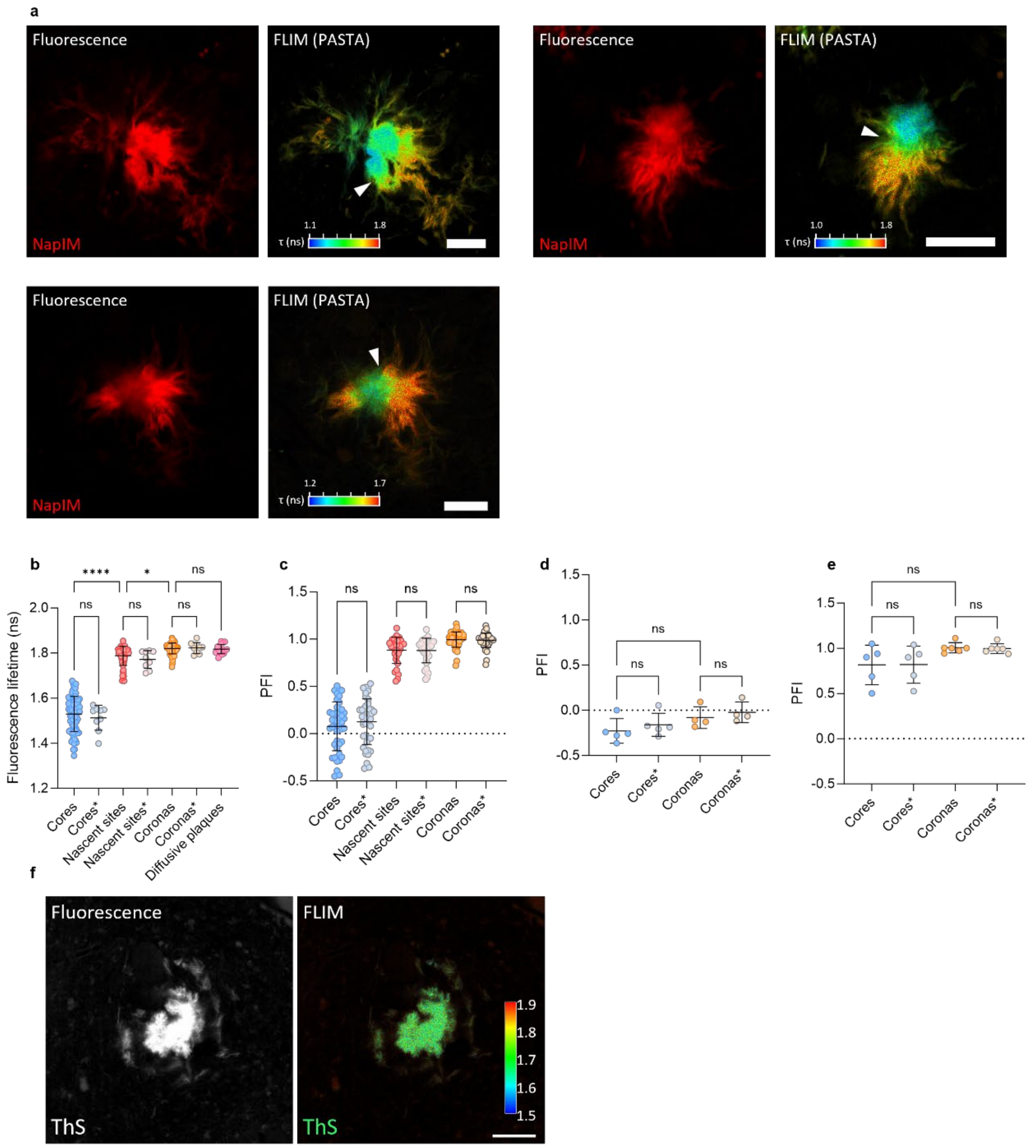

**Extended Data Fig. 5 | Discrimination of Aβ plaque heterogeneity by PASTA. a**, Representative fluorescence and lifetime (NapIM signal) images of Aβ plaque from a 12-month-old 5×FAD mouse labeled by NapIM. The transitional regions at subregion interfaces are indicated by white arrows. Scale bar, 10 μm. **b**, Fluorescence lifetime values of Aβ plaque subregions. Labels lacking asterisks indicate the full dataset: *n* = 95 plaques from *n* = 3 mice (12-month-old, 5×FAD); labels with asterisks indicate limited sampling from *n* = 9 classic cored plaques from *n* = 3 mice. **c–e**, PFI values of Aβ plaque subregions from classic cored plaques (**c**), "cold" classic cored plaques (**d**), and "hot" classic cored plaques (**e**). Labels lacking asterisks indicate the full dataset: *n* = 95 plaques from *n* = 3 mice (12-month-old, 5×FAD); labels with asterisks indicate PFI calculated from limited sampling, *n* = 9 classic cored plaques from *n* = 3 mice. **f**, Representative fluorescence (left) and lifetime (right) images of an Aβ plaque from a 12-month-old 5×FAD mouse labeled by ThS. Scale bar, 10 μm.

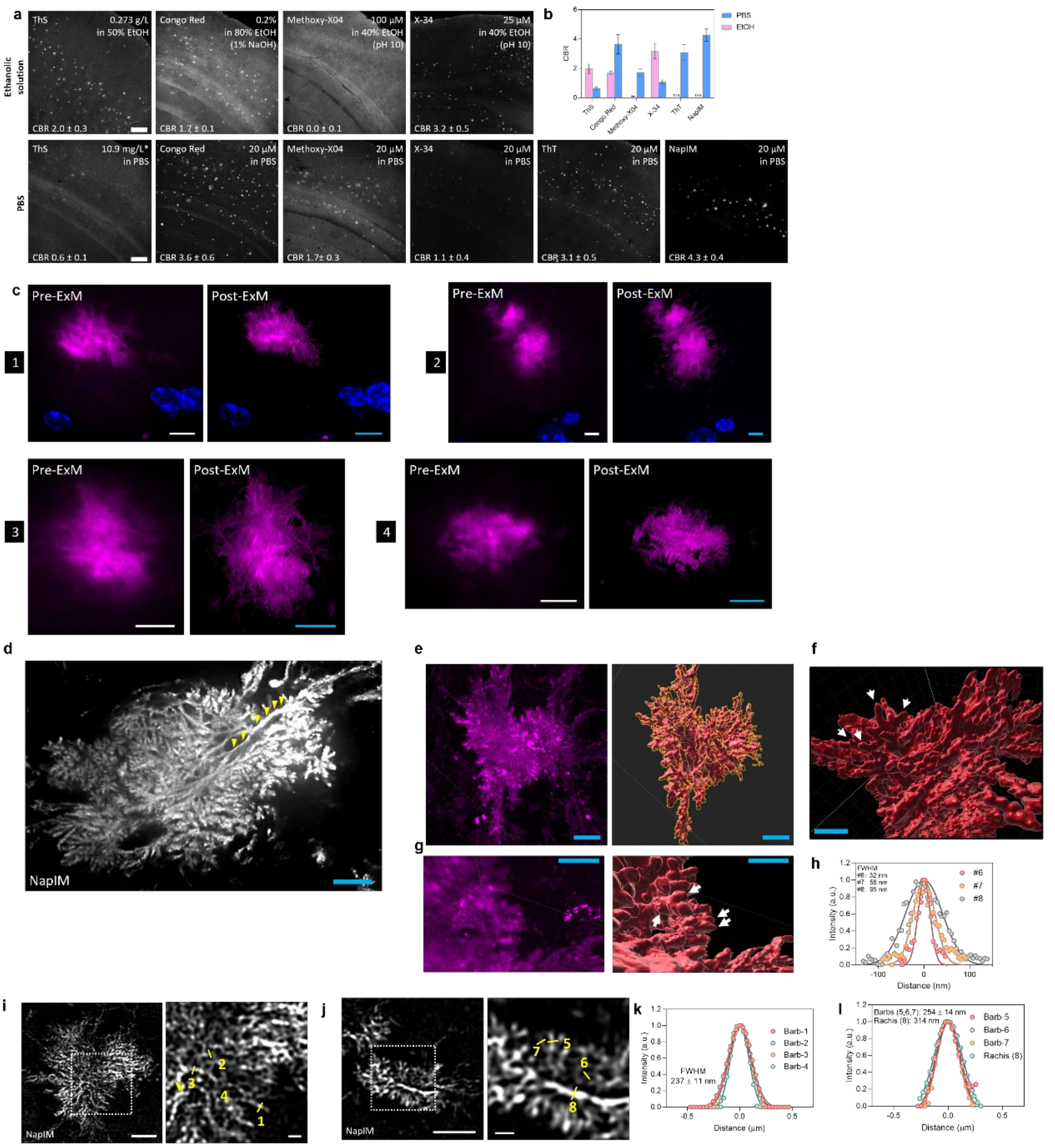

**Extended Data Fig. 6 | Visualization of Aβ plaques by NapIM on non-expanded tissue or using ExM. a,** Representative confocal images of Aβ plaques labeled by different probes from the same 12-month-old male 5×FAD mouse. Brain slices without any expansion treatments were treated with probes either in ethanolic solution based on reported protocols (ThS: [69]; Congo Red: [25, 70]; Methoxy-X04: [71]; X-34: [72]) or in 20 μM PBS solution. The average contrast-to-background ratios (CBRs) are indicated. Scale bar, 200 μm. Signals of Methoxy-X04, X-34, and ThT were measured on the 405 channel, ThS on the 488 channel, Congo Red on the 561 channel, and NapIM on the 647 channel. **b**, Quantifications of the CBRs for the images shown in **a**. Error bars, s.d., $n$ = 16. n/a, not available. *Since ThS is not chemically well defined, this concentration of ThS was used based on the reported structure [27] to roughly match the concentration of 20 μM. **c,** Representative pre- and post-expansion (5.6-fold) images of Aβ plaques from a 3-month-old 5×FAD mouse labeled by NapIM (magenta) to show the enhanced resolution. Scale bars, 5 μm in biological scales. **d,** A representative NapIM-labeled Aβ plaque from a 12-month-old 5×FAD mouse under ExM (4.5-fold) showing a distinctive main stem (indicated by yellow arrows). Scale bars, 2 µm in biological scale. **e**, The zoom-out view of a representative plaque under ExM (6.5-fold) from a 12-month-old 5×FAD mouse. Left, z-stack images of the plaque labeled by NapIM. Right, the 3D reconstruction of the plaque. **f–g**, Aβ branches were shown from the reconstruction. All scale bars, 1 µm in biological scale. **h**, Gaussian fit of the fluorescence profiles of barbs in Fig. 3l to yield FWHM. FWHM values of three barbs (labeled #6, #7, and #8 shown in Fig.3l) are indicated. **i–j**, Representative magnified SIM images of Aβ plaques from a 9-month-old 5×FAD mouse labeled by NapIM showing Aβ branches. Scale bars, 5 µm. Right, magnified views of plaque regions (boxes in left panel). Scale bar, 1 µm. **k–l**, Fluorescence intensity profiles marked by yellow lines with numbering in **i** and **j**, respectively, and Gaussian fits of the fluorescence profiles to yield full width at half maximum (FWHM). Symbols, experimental data; lines, Gaussian fit of the individual profiles.

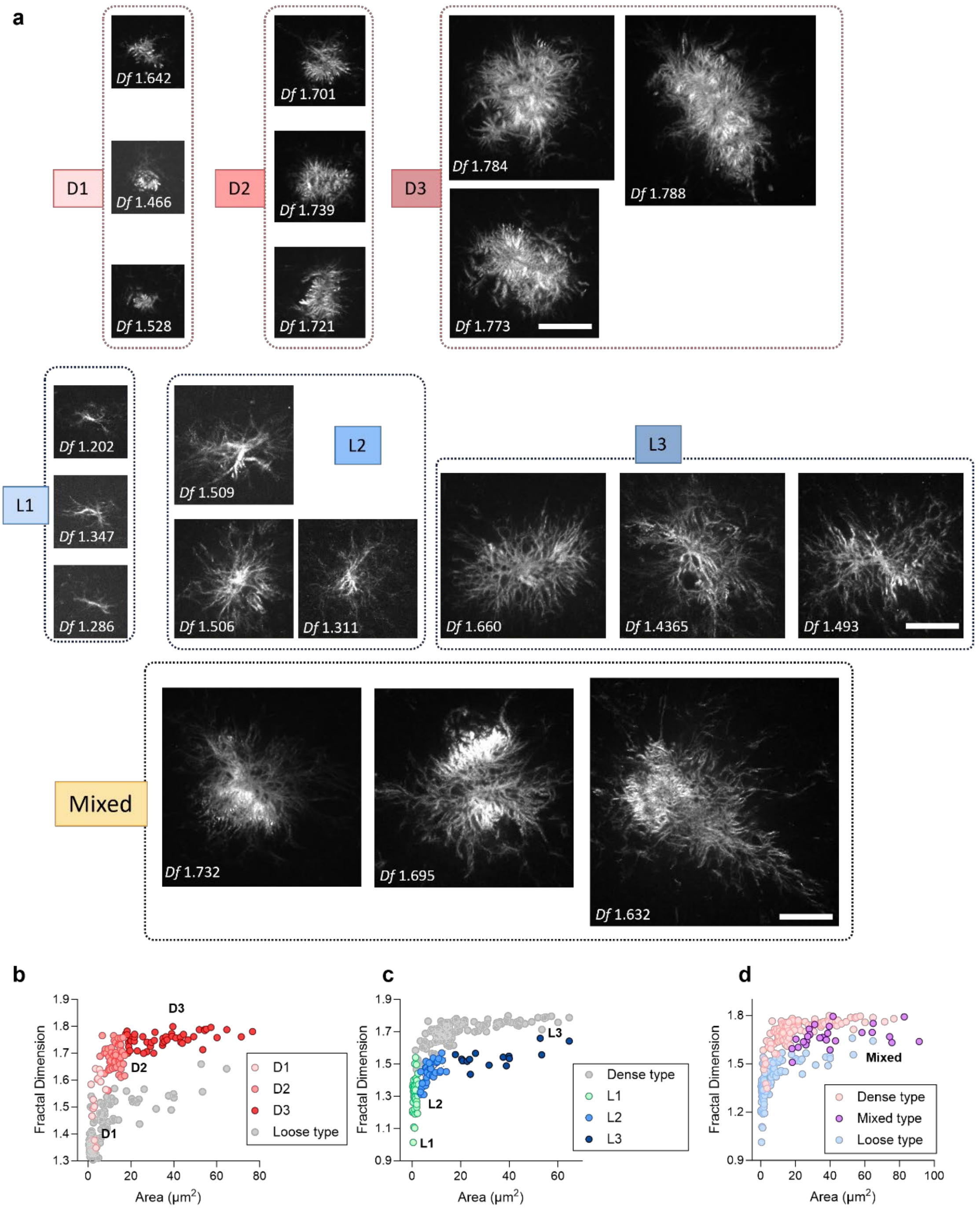

**Extended Data Fig. 7 | Aβ plaque phenotyping by PASTA. a**, Representative fluorescence images of dense type plaques with subtype D1, D2, and D3, loose type plaques with subtype L1, L2, and L3, and mixed type plaques, each showing 3 representative images. All images are shown at the same scale for ease of comparison, with all scale bars of 5 μm in biological scale. **b–j**, Plotting $D_f$ versus plaque areas shows distribution of subtypes of dense type (**b**), loose type (**c**) and mixed type (**d**).

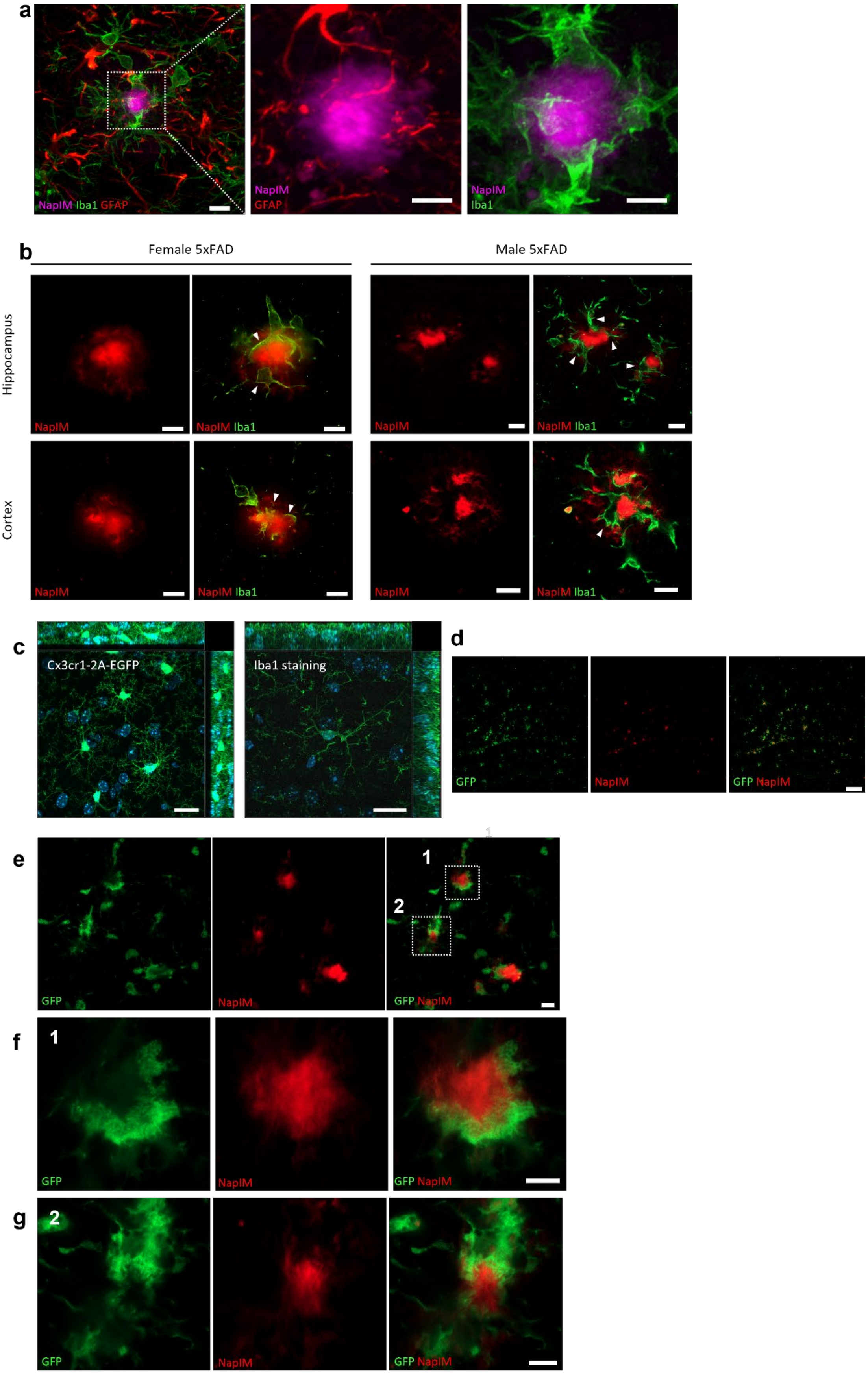

**Extended Data Fig. 8 | Visualization of interaction between microglia and Aβ plaques.**

**a,** Left, a representative confocal image of Aβ plaques (labeled by NapIM, magenta) and microglia by anti-Iba1 (green), astrocytes by anti-GFAP (red) from a 10-month-old 5×FAD mouse. Scale bar, 10 µm. Middle and right, magnified images of the dashed box showing more contact between Aβ and microglia than astrocytes. Scale bar, 5 µm. **b**, Representative confocal images of NapIM-labeled Aβ plaques (red) and microglia labeled by anti-Iba1 antibody (green) from male (10-month-old) and female (9-month-old) 5×FAD mice. White arrows indicate that microglial bodies fill the gap between dense cores and coronas. Scale bar, 10 µm. **c**, Cross-section projections of confocal z-stack images of microglia from CX3CR1-EGFP mice (1.5-month-old) or 5×FAD mouse (4-month-old) with anti-Iba1 staining. **d–e**, Representative confocal images of NapIM-labeled Aβ plaques and images with 10× (**d**) and 63× (**e**) magnification. **f–g**, Magnified regions of dashed box-1 (**f**) and box-2 (**g**) in **e**. Microglia were labeled by crossbreeding with CX3CR1-EGFP mice. Scale bars, 20 µm (**c**), 100 µm (**d**), 10 µm (**e**), 5 µm (**f, g**).

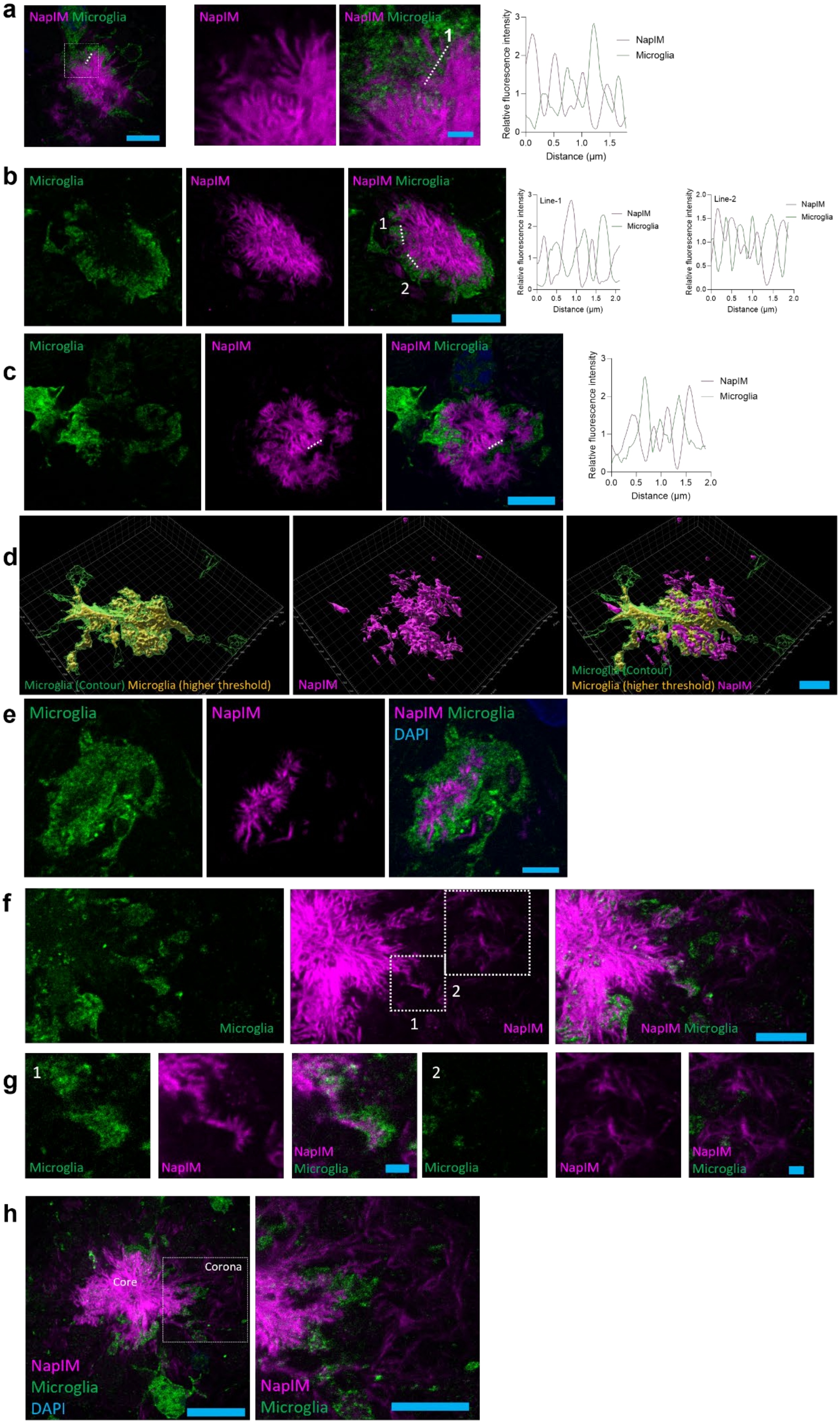
a
NapIM Microglia
NapIM
NapIM Microglia
1
Relative fluorescence intensity
Distance (μm)
NapIM
Microglia
b
Microglia
NapIM
NapIM Microglia
1
2
Line-1
Line-2
c
Microglia
NapIM
NapIM Microglia
d
Microglia (Contour) Microglia (higher threshold)
NapIM
Microglia (Contour)
Microglia (higher threshold) NapIM
e
Microglia
NapIM
NapIM Microglia
DAPI
f
Microglia
NapIM
NapIM Microglia
g
h
Core
Corona
NapIM
Microglia
DAPI

**Extended Data Fig. 9 | Visualization of microglial interaction with Aβ plaques using ExM.** **a**, Representative ExM images (6.0-fold) of Aβ plaques labeled by NapIM and microglia. Scale bar, 5 µm. Middle, magnified region of the dashed box. Scale bar, 1 µm. Right, line profile of fluorescence intensities across the interaction sites as shown by the dashed line. **b–c**, Representative ExM images (5.5-fold) of Aβ plaques labeled by NapIM and microglia. Scale bar, 5 µm. Right, line profiles of fluorescence intensities across the interaction sites are shown by dashed lines. **d**, 3D reconstruction of Aβ plaques (NapIM) and microglia under ExM (5.5-fold). Scale bar, 10 µm. **e**, Representative ExM images (6.0-fold) of Aβ plaques labeled by NapIM and microglia showing an Aβ plaque fully covered by microglia. Scale bar, 2 µm. **f**, Representative ExM images (4.5-fold) of Aβ plaques labeled by NapIM and microglia showed escaping Aβ branches and corona. Scale bar, 5 µm. **g**, Magnified regions of the dashed boxes in **f.** Scale bars, 1 µm. Scale bars are all on a biological scale. All Cx3cr1-2A-EGFP-labeled tissue slices were collected from 9-month-old Cx3cr1-2A-EGFP/5×FAD +/+ mice. **h**, Representative ExM images (4.5-fold) of Aβ plaques (labeled by NapIM) and microglia showing escaping Aβ branches and corona. Scale bar, 10 µm (biological scale). Right, magnified regions indicated by the white dashed box. Scale bar, 5 µm (biological scale).

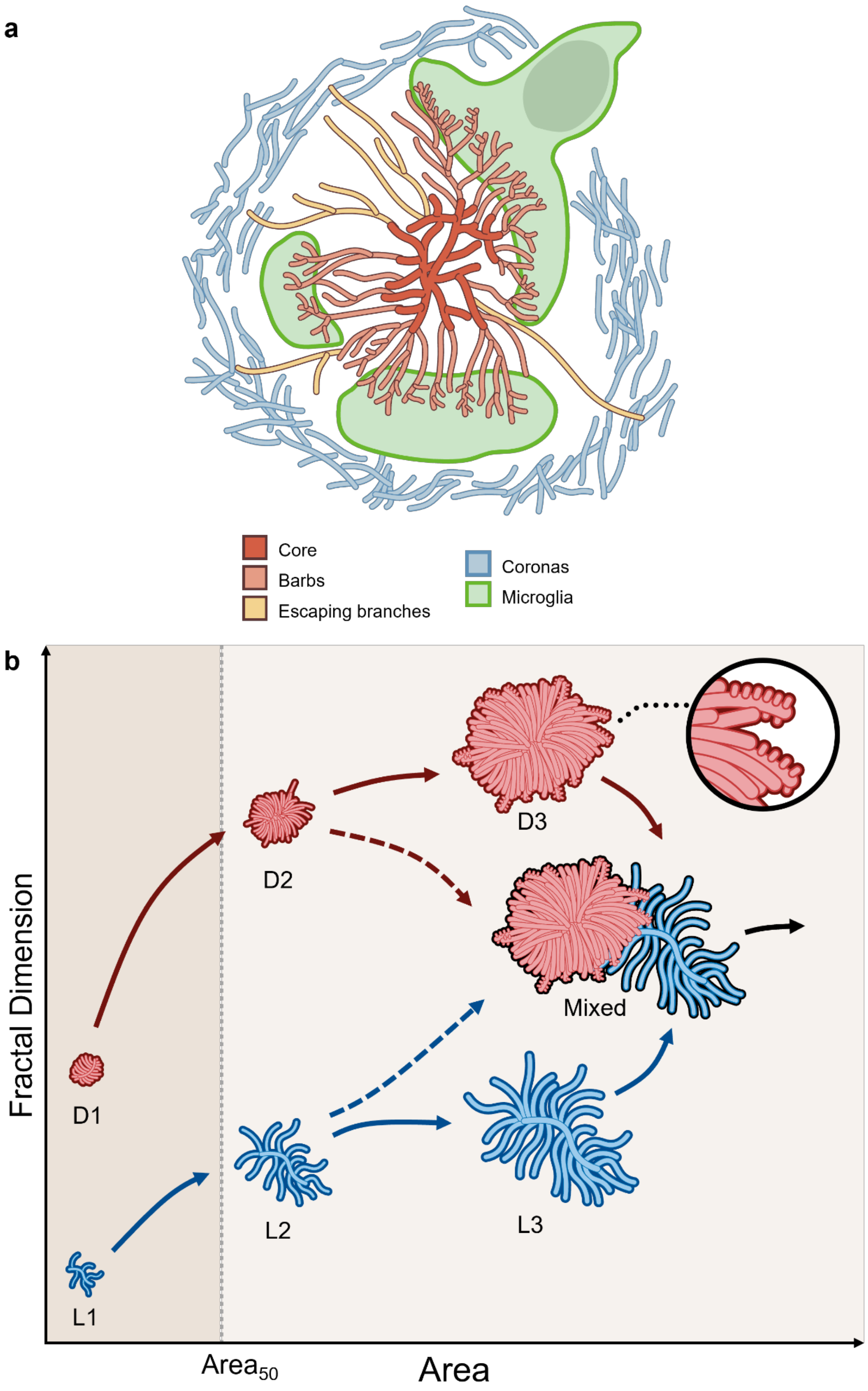

**Extended Data Fig. 10 | Proposed model of microglial interaction with Aβ plaques**

**and developmental trajectories in Aβ plaque evolution. a**, Schematic of Aβ plaque depicting smooth core, periphery, escaping branches, corona, microglial encapsulation, and rachis and barbs of pennaceous Aβ branches. **b**, Based on our study, we proposed a dual-lineage progression of Aβ plaques, originating from distinct early-stage precursors: the dense type (D1) and the loose type (L1). In this model, as plaques expand, they may follow lineage-specific growth patterns: the dense type undergoes extensive branching with increased Aβ branch density, whereas the loose type tends to elongate. Upon surpassing a critical size threshold, both lineages may transition into their respective intermediate phases (D2 and L2). Further maturation may lead to late-stage forms (D3 and L3); alternatively, inter-lineage convergence could occur, resulting in mixed-type plaques that incorporate features of both trajectories. Notably, our observations suggest distinct branch morphologies: dense lineages are characterized by spikey sprouts (barbs) projecting from branch surfaces, while loose lineages display predominantly smooth filaments.
